# Characterization of two *Martinezella* spp. species: *M. legmii* sp. nov. and *M. ivanii* sp. nov., from tropical legume trees in Costa Rica

**DOI:** 10.64898/2026.10.06.757023

**Authors:** Donifer Campos-Parra, Evelyn Artavia-Salazar, Bradd Mendoza-Guido, Mónica Vallejo-Arróliga, Ivonne Rodriguez-Ramirez, Keilor Rojas-Jimenez

**Affiliations:** Escuela de Biología, Universidad de Costa Rica, Sede Rodrigo Facio Brenes, 11501-2060 Montes de Oca, San José, Costa Rica; Instituto de Investigaciones en Salud (INISA), Universidad de Costa Rica, Sede Rodrigo Facio Brenes, 11501-2060 Montes de Oca, San José, Costa Rica

**Author notes:** **Corresponding author**: Keilor Rojas-Jimenez.

**Keywords:** Rhizobiaceae, Polyphasic taxonomic, *Arminella*, Nitrogen fixation, Fabaceae

## Abstract

Four rhizobial strains isolated from root nodules of tropical leguminous plants in Costa Rica were characterized. Strain LEGMi-135b was isolated from *Lonchocarpus felipei* root nodules, LEGMi-12c from *Cojoba arborea*, LEGMi-198b from *Calliandra haematocephala* and *Rhizobium altiplani* LEGMi-166a from *Mimosa pigra*. Phylogenetic analyses based on the 16S rRNA gene and four concatenated housekeeping genes (*recA, atpD, GATase* and *gyrB*) placed these strains within the genera *Martinezella* (LEGMi-12c, LEGMi-135b, and LEGMi-198b) and *Rhizobium* (LEGMi-166a). Genome-based analyses showed that strain LEGMi-135b exhibited an average nucleotide identity (ANI) value of 94% and a digital DNA-DNA hybridization (dDDH) value of 53.2% compared with its closest related species, *Martinezella hainanensis* CCBAU 57015^T^, supporting its classification as a distinct species. LEGMI-12c exhibited ANI and dDDH values of 98% and 87%, respectively, compared with *Martinezella hainanensis* CCBAU 57015^T^. LEGMi-198b showed the highest similarity to *Rhizobium cerradonense* CNPSo 3464, with ANI and dDDH values of 98% and 81.4%, respectively; however, this taxon has not been validly published. LEGMi-166a exhibited ANI and dDDH values of 100% and 99.3%, respectively, compared with *Rhizobium altiplani* BR 10423ᵀ, confirming its assignment to this species. Additionally, differentiation of strains LEGMi-135b and LEGMi-198b from their closest phylogenetic relatives was supported by phenotypic, physiological, and biochemical analyses. Based on the combined genomic, phenotypic, and biochemical evidence, we propose the establishment of the novel rhizobial species *Martinezella legmii* sp. nov., with strain LEGMi-135b designated as the type strain (LEGMi-135bᵀ), and *Martinezella ivanii* sp. nov., with strain LEGMi-198b designated as the type strain (LEGMi-198bᵀ). This study expands our understanding of rhizobial diversity associated with nitrogen-fixing symbioses, particularly among tropical leguminous plants from Costa Rica.

**Data summary:** The raw reads of the three *Martinezella* spp. and one *Rhizobium* sp. isolates have been deposited in GenBank under BioProject PRJNA1084876 and SRA accession SRP493729. The corresponding BioSample and genome accessions are: *Martinezella* sp. LEGMi-12c (SAMN44800730; GCF_054114145.1), LEGMi-135b (SAMN44800728; GCF_054114165.1), *Rhizobium* sp. LEGMi-166a (SAMN46966679; GCF_054114185.1), and *Martinezella* sp. LEGMi-198b (SAMN44800729; GCF_054114205.1). The 16S rRNA sequences are available under accessions PP439182, PP439210, PP439229, and PP439245, respectively.

## Introduction

The genera *Rhizobium* and *Martinezella* are part of the paraphyletic group known as rhizobia, which comprises various genera of Alphaproteobacteria and Betaproteobacteria classified within the orders Rhizobiales and Burkholderiales (1). *Martinezella* and *Rhizobium* belong to the family Rhizobiaceae, a group that formerly included approximately 38 genera, such as *Sinorhizobium*, *Mesorhizobium*, *Endobacterium*, *Pararhizobium*, *Pseudorhizobium*, *Shinella*, *Ciceribacter*, *Lentilitoribacter*, and *Agrobacterium,* among others (2). Recently, a study by Naranjo-Robayo et al. (3) proposed reclassifying 242 rhizobial strains into eight new genera, based on phylogenomic analyses and the requirement that genera be monophyletic (3).

Based on cpAAI (core-protein Amino Acid Identity) values falling below the <86.5% threshold and their phylogenetic placement, they reclassified the species *R. tumorigenes* 1078 and *R. tubonensis* CCBAU 85046 into the genus *Arminella* gen. nov. as *Arminella tumorigenes* and *Arminella tubonensis*, respectively. Likewise, they placed strains such as *R. calliandrae* CCGE524, *R. miluonensis* HAMBI 2971, *R. hainanensis* CCBAU 57015. *R. multihospitum* HAMBI 2975, *R. lusitana* P1 7, *R. rhizogenes* LBA9402, *R. jaguaris* CCGE525 and *R. dioscoreae* S 93 within the genus *Martinezella* gen. nov.

Nevertheless, species of the genera *Martinezella*, *Arminella*, and *Rhizobium* are Gram-negative soil bacteria belonging to the phylum Alphaproteobacteria (alpha-rhizobia) and the group of nitrogen-fixing bacteria known as rhizobia (4). *Martinezella* and *Rhizobium* are widely recognized for their symbiotic capacity with leguminous plants, with which they can form nodules and fix atmospheric nitrogen (5,6). These predominantly symbiotic species rely on symbiosis with host plants for nitrogen fixation and possess various genes that enable them to carry out these processes (7).

Nodulation capacity has generally been associated with *nod* genes, *nol* genes with the regulation (repression) of *nod* gene expression, and nitrogen fixation capacity with various genes, such as *nif* and *fix* genes (8,9). These genes can be physically located on plasmids and may therefore vary among different species, particularly due to their potential for horizontal gene transfer (7).

Symbiotic bacteria, even those belonging to the same species, can exhibit variation in the genetic makeup of the mobile genetic elements linked to their nodulation capacity, allowing them to be classified as symbiovars (7). A symbiovar is a specific adaptation of a species characterized by a gene combination that enables symbiosis with a particular plant species (7). Therefore, it is important not only to identify the *Martinezella*, *Arminella*, or *Rhizobium* species but also to identify differences in *nodC* genes; this helps elucidate their association with specific plant species, define their host specificity, and demonstrate whether these genes differ from those found in other symbiovars of the same species (7).

In our previous study, we determined *in silico* that strain LEGMi-135b corresponded to a novel species within the genus *Rhizobium*, whereas strain LEGMi-198b showed affinity with *Rhizobium cerradonense* Moura et al. 2023 -CNPSo 3464 (a taxon that has not yet been validly published), LEGMi-12c with *Rhizobium hainanensis* Chen et al. 1997 - CCBAU 57015^T^, and LEGMi-166a with *Rhizobium altiplani* Baraúna et al. 2016 - BR 10423ᵀ (10–12). However, following the analysis by Naranjo-Robayo et al. (13), it became necessary to update the taxonomic affiliations of those strains in light of the newly described genera *Martinezella* and *Arminella*.

In the present study, we phenotypically characterized two novel species, *Martinezella legmii* sp. nov. (LEGMi-135b) and *Martinezella ivanii* sp. nov. (LEGMi-198b), as well as two additional isolates, LEGMi-12c and LEGMi-166a. Furthermore, using phylogenomic and core-protein Amino Acid Identity (cpAAI) analyses, we evaluated and clarified the taxonomic relationships of our strains with closely related species belonging to the genera *Rhizobium*, *Martinezella*, and *Arminella*.

## METHODS

### Isolation and ecology

Strain LEGMi-135b was isolated from *Lonchocarpus felipei* nodules, LEGMi-12c from *Cojoba arborea*, LEGMi-198b from *Calliandra haematocephala*, and *Rhizobium altiplani* LEGMi-166a from *Mimosa pigra*, four tree legume species from the Costa Rican tropics (14). The nodules were harvested and surface-sterilized, and the bacteria were isolated following the protocol previously described by Rojas-Jimenez et al. (2026) (14–16). The nodules were washed and disinfected using sequential treatments with 70% ethanol and 2% sodium hypochlorite, followed by rinsing with distilled water. They were macerated in sterile phosphate-buffered saline (PBS, pH 7.4) to release the bacterial content, and the resulting suspension was subjected to serial dilutions in PY broth (5 gL^−1^ peptone, 1 gL^−1^ anhydrous dextrose, 0.5 gL^−1^ dipotassium phosphate, 0.2 gL^−1^ magnesium sulfate, and 0.1 gL^−1^ sodium chloride). Aliquots of 50 μL from each dilution were plated on PY medium supplemented with cycloheximide (40 mgL^−1^) and incubated at 28°C for three weeks. Individual colonies were purified by re-streaking onto the same medium. Finally, pure bacterial colonies were preserved in PY broth with 20% glycerol and stored at -80°C in the collection of the Laboratory of Genetics and Ecology of Microorganisms (LEGMi) at the School of Biology, University of Costa Rica (14).

### Genome-Based Relatedness and Phylogenomic Analyses

To supplement the sequence data obtained in our previous study, we re-cultured the isolates on PYA and extracted DNA using the DNeasy PowerSoil Kit (Qiagen), following the manufacturer’s instructions. Whole-genome sequencing was performed by Novogene (CA, USA) using the Illumina NovaSeq PE150 platform. Consequently, by combining these with the data from our initial study, we obtained two sets of Illumina reads for strains LEGMi-12c, LEGMi-135b, and LEGMi-166a. Additionally, for strain LEGMi-135b, we had sequence data previously generated using PacBio technology.

Raw Illumina reads were processed with fastp v0.20.1 (default parameters, Q30 threshold), and the resulting reads were assembled with SPAdes (v.3.15.4) using k-mers of 21, 33, 55, and 77 (17,18). Additionally, strain LEGMi-135b was assembled using the hybrid Illumina–PacBio approach implemented in SPAdes via the hybridSPAdes pipeline (19). The hybridSPAdes pipeline integrates long-read information to resolve repeats and close gaps in short-read assemblies using the exSPAnder module (19). The assembly was performed using k-mer sizes of 33, 55, 77, 99, and 127. Contigs <1,000 bp were removed from all assemblies using seqtk (20). Assembly statistics were generated with Quast, while genome completeness and contamination were evaluated using CheckM2 v1.0.1 (21,22). We used eggNOG-mapper v2.1.6, NCBI Prokaryotic Genome Annotation Pipeline (PGAP), and RAST tools for genome annotation (23–25).

We manually compiled a list of GenBank accession numbers for species closely related to our isolates, based on the results of a literature review, an examination of genomes available in GenBank, and whole-genome comparisons performed using the Type Strain Genome Server (TYGS), based on digital DNA-DNA hybridization (dDDH4), and the Genome Taxonomy Database Toolkit (GTDB-Tk) (26,27).

We used the Genome-to-Genome Distance Calculator 3.0 tool (formula 2, d4) to estimate digital DNA-DNA hybridization (dDDH) with the species most closely related to our isolates (28). Additionally, we calculated average nucleotide identity (ANI) values using pyANI (29).

For the phylogenetic analysis of the core genome, we first performed a broad analysis including our strains and 87 bacterial species from the genera *Rhizobium*, *Martinezella*, and *Arminella*. We used GenFlow, an in-house pipeline that retrieves user-specified reference genomes from GenBank (NCBI) and processes them using Anvi’o (v.7.1) (30,31). These taxa were identified through the List of Prokaryotic names with Standing in Nomenclature (LPSN) and included genomes available in GenBank that met the inclusion criteria of being reference genomes and, preferably, being derived from type material (1,32). We also included genomes from species that, according to the LPSN, hold a “not validly published” nomenclatural status but serve as reference genomes not associated with type material. Within the pipeline, coding sequences (CDS) were annotated using Prodigal, and shared core single-copy genes (SCCGs) were identified and extracted. These sequences were aligned using MAFFT v7.397, resulting in a concatenated alignment comprising 651 single-copy orthologous gene sequences and a total of 59,241 genes. The phylogenomic tree was subsequently constructed using FastTree with the JTT+CAT model and visualized using iTOL v6 (33) (Fig. S1).

Subsequently, to obtain a more focused view of the phylogenetic relationships within *Bradyrhizobium*, we performed a second core-genome analysis including the available reference genomes of the genus. Using GenFlow, coding sequences (CDS) were annotated with Prodigal, and shared core single-copy genes (SCCGs) were identified and extracted. The sequences were aligned using MAFFT v7.397, resulting in concatenated amino acid sequences derived from 1,555 single-copy core gene clusters containing a total of 57,535 genes. The resulting phylogenomic tree was constructed using FastTree with the JTT+CAT model and visualized using iTOL v6. Additionally, we employed the pipeline proposed by Naranjo-Robayo et al. (13) to calculate the core-proteome Average Amino Acid Identity (cpAAI) between our strains and the reference genomes of the genera *Rhizobium*, *Martinezella*, and *Arminella*, in order to determine the correct taxonomic designation of our study strains at the genus level (13). Phylogenetic inference was performed based on the concatenated alignment of the 170 marker proteins originally established by Kuzmanović et al. (34), using IQ-TREE v2.2.0 (35). A threshold of ∼86.5% was used to delimit genera, as adjusted by Naranjo-Robayo et al. (13).

In the first stage, the most suitable amino acid substitution model was automatically selected using the ModelFinder option (-m MF), restricting the search to the LG model family (--mset LG) and selecting based on the Bayesian Information Criterion (BIC) (13). Subsequently, a Maximum Likelihood (ML) tree was reconstructed using all previously selected reference genomes from *Rhizobium*, *Martinezella*, and *Arminella* under the best-fit model LG+F+R5 (Fig. S2). Next, a phylogenetic tree was inferred for our strains alongside their closest phylogenetic relatives, as identified in the previous analysis, using the LG+F+R5 model. Branch robustness was assessed via 1,000 ultrafast bootstrap replicates (-J 1000) and 1,000 replicates of the approximate likelihood-ratio test (SH-aLRT; --alrt 1000). The resulting phylogenetic trees and heatmaps of cpAAi values were visualized using iTOL v6. The corresponding sequence alignments and phylogenetic trees in Newick format are provided as supplementary material

### *16s rRNA, nodC,* House Keeping genes phylogeny

16S rRNA sequences and housekeeping genes (*recA*, *atpD*, *GATase*, and *gyrB*) were extracted from complete genomes available in GenBank (NCBI) using the “anvi-get-sequences-for-hmm-hits” tool in Anvi’o v7.1, with gene detection performed via HMMER v3.3.2 (31,36).

To identify the *nodC* gene, reference sequences were initially obtained from UniProt and curated by removing those containing ambiguous or non-standard amino acids (B, J, O, U, X, and Z) (37). Subsequently, the filtered sequences were aligned using MAFFT v.7.397; the alignment was refined with trimAl using the “automated1” mode, and the resulting alignment was used to construct a Hidden Markov Model (HMM) with HMMER (hmmbuild) (36,38,39).

Predicted amino acid sequences were extracted from each contig database generated with Anvi’o using the “anvi-get-sequences-for-gene-calls” tool, yielding a set of proteomes comprising the genomes assembled for the study and the reference genomes of symbiovars, specifically those with complete genomes, compiled by Martinez-Romero et al (7). Symbiovars lacking complete genomes, possessing only partial *nodC* sequences, were excluded from this analysis.

These sequences were used to search for and identify *nodC* homologs using the constructed HMM model. The identification of *nodC* orthologs within each proteome was performed using “hmmsearch” from the HMMER package, utilizing the previously constructed HMM profile. For each genome, the match with the highest bit score was retained. In cases where two or more matches yielded the same bit score, the match with the lowest E-value was selected, as this represented the evidence with greater statistical significance.

The sequence corresponding to the best alignment was extracted from the proteome using SeqKit v0.7.1 (40), yielding a single putative *nodC* ortholog per genome. To confirm that the putative orthologous sequence corresponded to *nodC*, a BLASTp analysis was performed. Sequences that did not exhibit identity consistent with *nodC* proteins (chitooligosaccharide synthase NodC, N-acetylglucosaminyltransferase, or nodulation protein C) were discarded (41).

Finally, the 16S rRNA, *nodC*, and housekeeping gene sequences were aligned separately using MAFFT v7.397 (38). TrimAl (in “automated1” mode) was used to automatically remove spurious sequences or misaligned regions from the housekeeping, 16s rRNA and *nodC* gene sequences (39). Maximum-likelihood phylogenetic trees were inferred for the 16S rRNA, housekeeping, and *nodC* genes using IQ-TREE v2.2.0 with the ModelFinder Plus option (-m MFP), which determines the best-fit substitution model based on the Bayesian Information Criterion (BIC) (35). Trees were inferred using the TIM3+F+I+G4 model for the 16S rRNA gene, the Q.insect+F+I+G4 model for housekeeping genes, and the Q.pfam+F+R4 model for *nodC*; all inferences included 1,000 bootstrap replicates. Additionally, to identify the symbiovar most closely related to the strains, a percent identity matrix was calculated using Clustal Omega, applying a 95% threshold for symbiovar delimitation, and a heatmap was generated (7,42). Finally, all phylogenetic trees were visualized using iTOL (v.6.9.1) (33). The corresponding sequence alignments and phylogenetic trees in Newick format are provided as supplementary material.

### Nodulation test

We conducted in vivo nodulation tests using *Phaseolus vulgaris* plants for strains LEGMi-12c (two replicates), LEGMi-135b (four replicates), LEGMi-166a (two replicates) and LEGMi-198b (four replicates). Seeds of *P. vulgaris* cv. Cabécar were selected and washed twice with distilled water; they were then immersed in 70% ethanol for 4 minutes. The ethanol was decanted, and a 20% (v/v) sodium hypochlorite solution containing 2 drops of Tween 20 was added, followed by agitation for 15 minutes (43). Then, the seeds were washed three times with sterile distilled water, and 70 mL of a 0.01% Benomyl solution was added with agitation for 4 minutes. Afterward, the solution was decanted, and the seeds were washed three times with sterile distilled water. The seeds were dried using sterile paper and placed (2 to 3 seeds per container) in jars containing germination medium (4.4 gL^−1^ MS salts, 0.1 gL^−1^ myo-inositol, 1000 µL thiamine, 30 gL^−1^ sucrose, and 6 gL^−1^ Phytagel); the jars were sealed with paraffin and incubated at 25°C in the dark for 7 days (44).

Seedlings were selected based on complete germination, developed radicles and plumules, similar radicle lengths (2–5 cm), and the visible absence of cotyledon damage or bacterial/fungal contamination. These were transferred to a semi-hydroponic system using modified Leonard jars containing a 1:2 perlite-vermiculite substrate mixture in the upper section and nitrogen-free Fahraeus (NF) medium (15 gL^−1^ disodium phosphate, 10 gL^−1^ monopotassium phosphate, 10 gL^−1^ calcium chloride, 12 gL^−1^ magnesium sulfate, 0.5 gL^−1^ iron(III) citrate, 2.86 gL^−1^ boric acid, 2.003 gL^−1^ manganese(II) sulfate, 0.22 gL^−1^ zinc sulfate, 0.08 gL^−1^ copper(II) sulfate, and 0.08 gL^−1^ sodium molybdate) in the lower section (45). They were grown in a temperature controlled environment at 23 °C.

Six days later, they were inoculated with 1 mL of fresh culture of strains LEGMi-12c, LEGMi-135b, LEGMi-166a and LEGMi-198b in YM broth at an absorbance between 0.2 and 0.4 (OD: 600 nm). Then, they were inoculated a second time at six days and a third time at twenty days. The plants were harvested 40 days after transfer to Leonard jars; the roots were washed with sterile distilled water and examined for the presence of nodules. The nodules found were preserved in 1% PBS and stored at -80°C.

### Physiology and chemotaxonomy

We cultured the study strains on the following media: LB (10 gL^−1^ tryptone, 5 gL^−1^ yeast extract, 10 gL^−1^ sodium chloride (NaCl), and 15 gL^−1^ agar), PY (5 gL^−1^ peptone, 1 gL^−1^ anhydrous dextrose, 4 gL^−1^ mannitol, 0.5 gL^−1^ dipotassium phosphate, 0.2 gL^−1^ magnesium sulfate, 0.1 gL^−1^ sodium chloride, and 15 gL^−1^ agar), YMA (1 gL^−1^ yeast extract, 4 gL^−1^ mannitol, 0.5 gL^−1^ dipotassium phosphate, 0.2 gL^−1^ magnesium sulfate, 0.1 gL^−1^ sodium chloride, and 15gL^−1^ agar), PGA (10 gL^−1^ peptone, 5 gL^−1^ anhydrous dextrose, 5 gL^−1^ sodium chloride, and 15 gL^−1^ agar), MacConkey, and Mueller-Hinton. Additionally, strain growth was tested on YMA medium supplemented with Congo red and bromothymol blue indicators, and on PGA medium with 0.1% bromocresol purple. Colony morphology was observed on PY+ medium. To determine shape and size, and to classify the isolated strains based on cell wall structure, we performed Gram staining and observed the samples using light microscopy at 100x magnification and transmission electron microscopy (TEM) at 8000x magnification.

We determined the effect of temperature on growth by inoculating the isolated strains onto YMA medium plates, incubating them at various temperatures (20°C, 25°C, 30°C, and 35°C), and examining them after 48 hours. Salt tolerance was determined by inoculating the strains into PYA medium supplemented with 1% and 2% NaCl (w/v) and assessing growth after 48 hours. We used YMA medium supplemented with bromothymol blue (2.5%) in plates to qualitatively determine medium acidification or alkalinization resulting from bacterial growth, based on the color change of the medium.

We determined the isolates’ growth capacity at different pH levels by placing them in YM broth adjusted to various pH values (2, 4, 5, 7, 9, and 11). pH adjustment was achieved by adding varying concentrations of HCl and NaOH to 60 mL of YM broth. Subsequently, we inoculated the medium with 500 µL of the isolated strain, previously grown in YM broth to an absorbance (OD: 600 nm) between 0.155 and 0.400, and performed an initial absorbance measurement; finally, after 24 hours of incubation with agitation, we measured the absorbance again.

The bioMérieux VITEK® 2, an automated identification system using the Gram-Negative (GN) card, which comprises 47 colorimetric biochemical tests, was used to assess carbon source utilization, inhibition, resistance, and enzymatic activity of the isolates (46,47). The GN card enables the taxonomic identification of at least 159 Gram-negative bacterial taxa, including *Rhizobium radiobacter* (*47,48*). Samples were processed according to the manufacturer’s instructions: a suspension of isolated colonies from the culture was prepared in 3.0 mL of sterile saline (0.45–0.50% NaCl, pH 4.5–7.0), and the cell density was adjusted to a McFarland turbidity standard of 0.50–0.63. Finally, the bacterial suspension and the GN card were loaded into the VITEK® 2 system for automated processing (47).

Antibiotic susceptibility of the study strains was determined using an antibiogram test; susceptibility to ampicillin (AMP), streptomycin (S), neomycin (N), cefuroxime (CXM), and vancomycin (VRE) was evaluated. Five colonies of each strain were suspended in a tube containing 9 mL of YM broth until an optical density (OD: 600 nm) of 0.090 to 0.130 (equivalent to a 0.5 McFarland standard) was reached at room temperature. This suspension was used to streak Mueller-Hinton agar plates in perpendicular and diagonal patterns to ensure the plate was fully covered with the bacteria. Antibiotic discs were then placed on the plates, and the cultures were incubated at 28°C for 24 hours. Finally, the appearance of an inhibition halo was qualitatively interpreted as indicating the strain’s susceptibility to the specific antibiotic.

## RESULTS AND DISCUSSION

### Genome-Based Relatedness and Phylogenomic Analyses

LEGMi-166a presented the largest genome, followed by LEGMi-12c, LEGMi-198b, and LEGMi-135b; this same order was observed regarding the number of genes (Table 1). All genomes showed completeness close to or equal to 100% and contamination close to 0%. LEGMi-198b was the assembly with the lowest number of contigs, followed by LEGMi-135b, LEGMi-166a, and LEGMi-12c. The G+C content in all strains was close to 60%. Additionally, all isolated strains contain *fix*, *nif*, *nod*, *nol*, and *ntr* genes, suggesting *in silico* potential for nodulation and nitrogen fixation (Table 1, Table S1). The complete PGAP annotation is publicly available online, while the EggNOG-mapper and RAST annotation results are provided as supplementary materials.

**Table 1.** Genome features of the three *Martinezella* spp. and the *Rhizobium* sp. isolates.

| Strain characteristic | Data for: |  |  |  |
| --- | --- | --- | --- | --- |
|  | *Martinezella* sp. -LEGMi-12c | *Martinezella* sp. -LEGMi-135b | *Rhizobium* sp. -LEGMi-166a | *Martinezella* sp. -LEGMi-198b |
| Legume host species | *Cojoba arborea* | *Lonchocarpus felipei* | *Mimosa pigra* | *Calliandra haematocephala* |
| No. of read (post QC) | 8,188,458 | 10,503,224 | 9,500,990 | 8,669,270 |
| No. of contigs | 119 | 33 | 107 | 22 |
| No. of genes | 6454 | 6192 | 7,464 | 6357 |
| Genome length (bp) | 6,846,952 | 6,620,416 | 7,565,831 | 6,828,554 |
| Coverage (x) | 179.39 | 237.97 | 210.30 | 190.43 |
| N50 (bp) | 109,404 | 571,440 | 185,397 | 817,417 |
| GC content (%) | 59.82 | 59.77 | 59.48 | 59.53 |
| Completeness (%) | 99.94 | 99.98 | 100 | 100 |
| Contamination (%) | 0.47 | 0.15 | 1.69 | 0.18 |
| Closest 16S rRNA match – accession | *Rhizobium* sp. strain IITA-TZ190 - OM909439 | *Rhizobium* sp. strain IITA-TZ190 - OM909439 | *Rhizobium altiplani* strain BR10893 - OM144531 | *Rhizobium rhizogenes* strain SQL8 -MN480505 |
| Nodulation genes) | *nodL*, *nodN*, *nodQ*, *nodT* and *nodW* | *nodL*, *nodN*, *nodQ*, *nodT* and *nodW* | *nodA*, *nodB*, *nodC*, *nodC*, *nodD*, *nodH*, *nodI*, *nodJ*, *nodN*, *nodQ*, *nodS*, *nodU* and *nodZ* | *nodA*, *nodB*, *nodC*, *nodD*, *nodD1*, *nodI*, *nodJ*, *nodL*, *nodN*, *nodQ*, *nodS*, *nodT*, *nodU* and *noIO* |
| Nitrogen fixation genes | *fixH*, *fixK* and *nifU* | *fixK* and *nifU* | *fixA*, *fixB*, *fixC*, *fixG*, *fixH*, *fixI*, *fixJ*, *fixK*, *fixQ*, *fixR*, *fixS*, *fixU*, *fixX*, *nifA*, *nifB*, *nifD*, *nifE*, *nifH*, *nifK*, *nifN*, *nifQ*, *nifS*, *nifU*, *nifW*, *nifX* and *nifZ* | *fixA*, *fixB*, *fixC*, *fixG*, *fixH*, *fixI*, *fixJ*, *fixK*, *fixL*, *fixQ*, *fixS*, *fixU*, *fixX*, *nifA*, *nifB*, *nifD*, *nifE*, *nifH*, *nifK*, *nifN*, *nifQ*, *nifS*, *nifT*, *nifU*, *nifX* and *nifZ* |
| Regulatory genes | *noIR*, *ntrB*, *ntrC*, *ntrX* and *ntrY* | *noIR*, *ntrB*, *ntrC*, *ntrX* and *ntrY* | *noIR*, *ntrB*, *ntrC*, *ntrX* and *ntrY* | *noIR*, *noIR*, *ntrB*, *ntrC*, *ntrX* and *ntrY* |

The strains isolated in the study were taxonomically classified as belonging to the genera *Martinezella* spp. (LEGMi-12c, LEGMi-135b) and *Rhizobium* spp. (LEGMi-198b and LEGMi-166a) according to GTDB-tk and TYGS (Table S2). GTDB-tk was unable to identify a FastANI reference for strain LEGMi-135b. After compiling a list of 87 reference genomes, including the genera *Arminella*, *Martinezella*, and *Rhizobium*, available in GenBank and reconstructing the core-genome phylogenomic tree, bootstrap support values for key nodes were found to be 100% in all cases (Fig. 1). Strain LEGMi-12c is closely related to *Martinezella hainanensis*, whereas strain LEGMi-135b, forms a branch independent of LEGMi-12c and *Martinezella hainanensis*. LEGMi-198b forms a clade with *Rhizobium* sp. NXC24 and *Rhizobium cerradonense* CNPSo 3464, neither of which is validly published according to LPSN, within the *Martinezella* clade (*13*). Meanwhile, LEGMi-166a is phylogenetically related to *R. altiplani* BR 10423T and falls within the “leguminosarum-etli” clade (13). Analysis of concatenated housekeeping gene sequences (*recA*, *atpD*, *GATase*, and *gyrB*) confirms the phylogenetic relationships observed in the core-genome phylogenomic tree between the study strains and their closest relatives (Fig. 2).

**Fig. 1.**
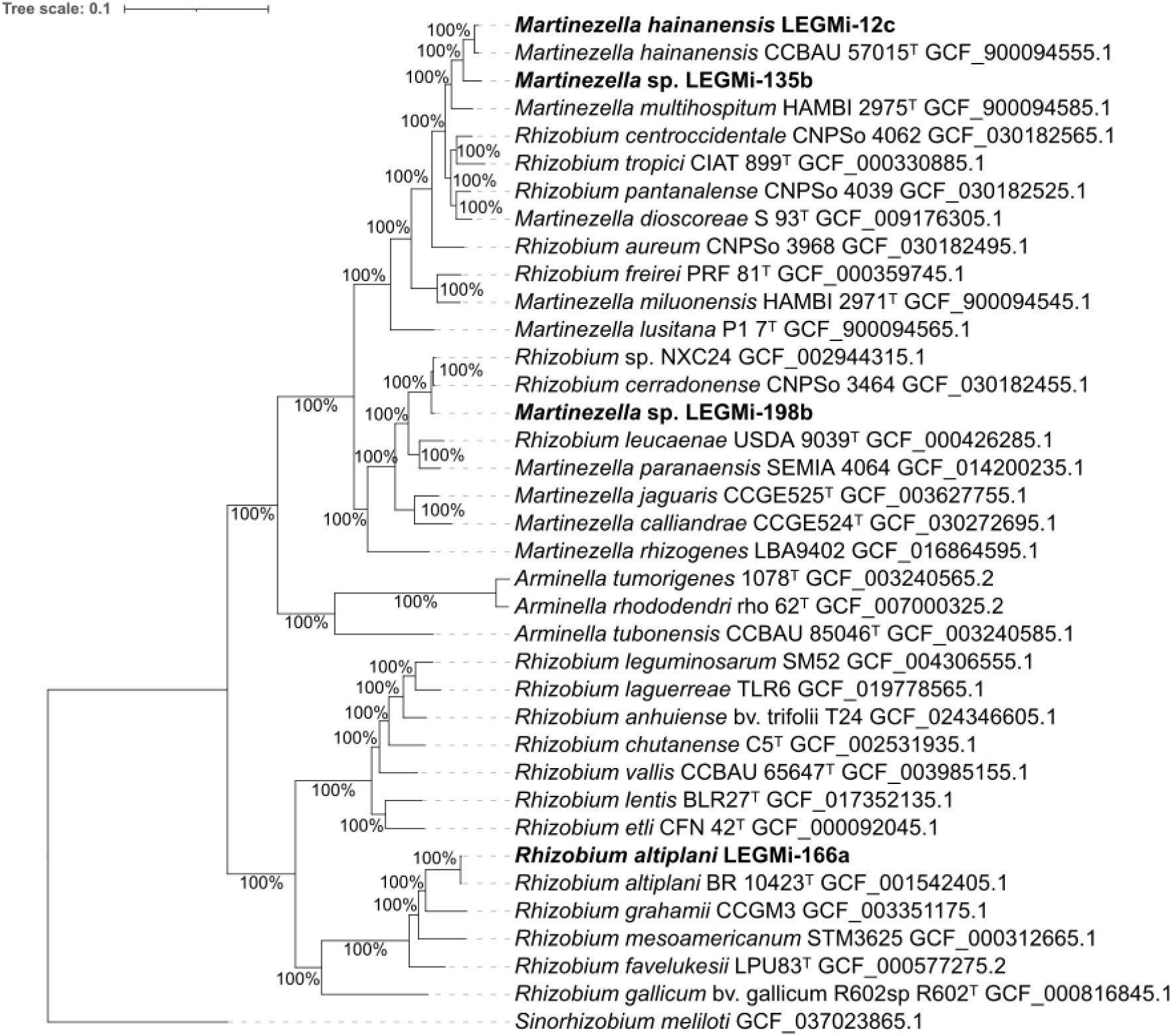
Phylogenomic tree inferred from the core genome of 37 bacterial genomes, comprising 1,555 single-copy orthologous gene sequences and a total of 57,535 genes. Phylogenetic reconstruction was performed using FastTree under the JTT-CAT substitution model with 1,000 bootstrap replicates. Reference genomes were manually selected from the GenBank database, giving priority to type strains and including the closest available *Rhizobium*, *Martinezella* and *Arminella* genomes identified through TYGS analyses, published studies, and, whenever available, reference genomes associated with type material. Genome accession numbers are indicated alongside each strain name. Type strains are denoted by a superscript “T”, while bootstrap support values are shown at the corresponding nodes. The three putative *Martinezella* and the *Rhizobium* species identified in this study are highlighted in bold. The genome of *Sinorhizobium meliloti* was used to root the tree. Tree visualization and annotation were carried out using iTOL v6.

**Fig. 2.**
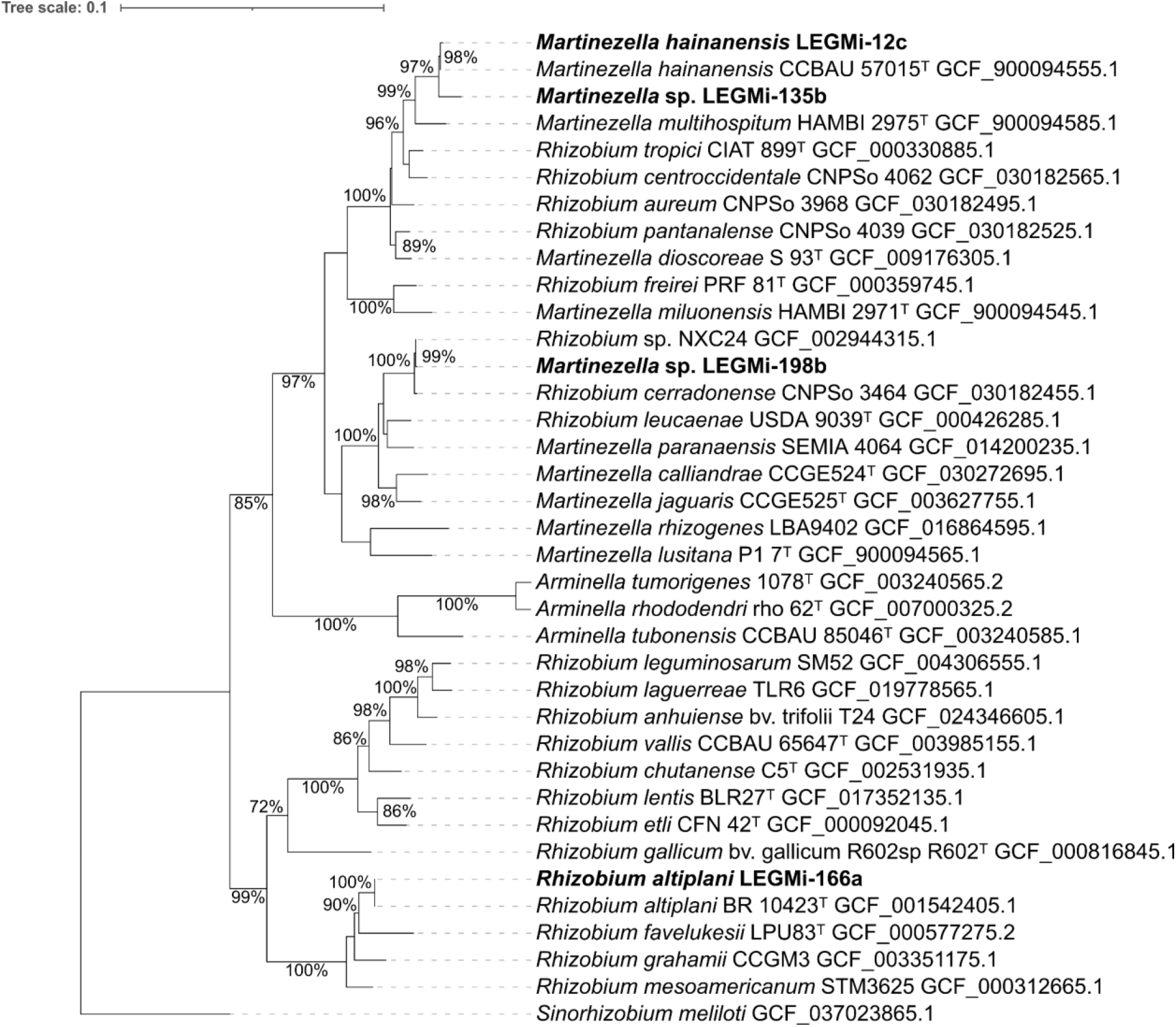
Maximum-likelihood phylogenetic tree inferred from the concatenated housekeeping genes (*recA, atpD, GATase* and *gyrB*). The tree was reconstructed with IQ-TREE v2.2.0 under the Q.insect+F+I+G4 substitution model. A total of 37 genomes from the genera *Rhizobium*, *Martinezella*, and *Arminella*, corresponding to the closest relatives of the isolates characterized in this study, were included in the analysis. Genome accession numbers are provided next to each strain name, and type strains are indicated by the superscript “T”. Bootstrap support values ≥70% are displayed at the corresponding nodes. The three putative *Martinezella* strains and the *Rhizobium* species identified in this study are shown in bold. The tree was rooted using the genome of *Sinorhizobium meliloti* as the outgroup. Visualization and annotation were performed in iTOL v6.

The cpAAI analysis of the genomes showed that strains LEGMi-12c, LEGMi-135b, and LEGMi-198b exhibit values above the proposed threshold for genus delimitation (cpAAI ≥ 86.5%) when compared to the species comprising the *Martinezella* genus cluster described by Naranjo-Robayo et al. (13) (Fig. 3, Fig. S2). In contrast, the cpAAI values obtained against species of the genera *Arminella* and *Rhizobium* were below this threshold, indicating lower genetic affinity with these genera. Strain LEGMi-166a belongs to the cluster formed by bacteria of the genus *Rhizobium* and exhibits cpAAI values above the genus delimitation threshold with them.

**Fig. 3.**
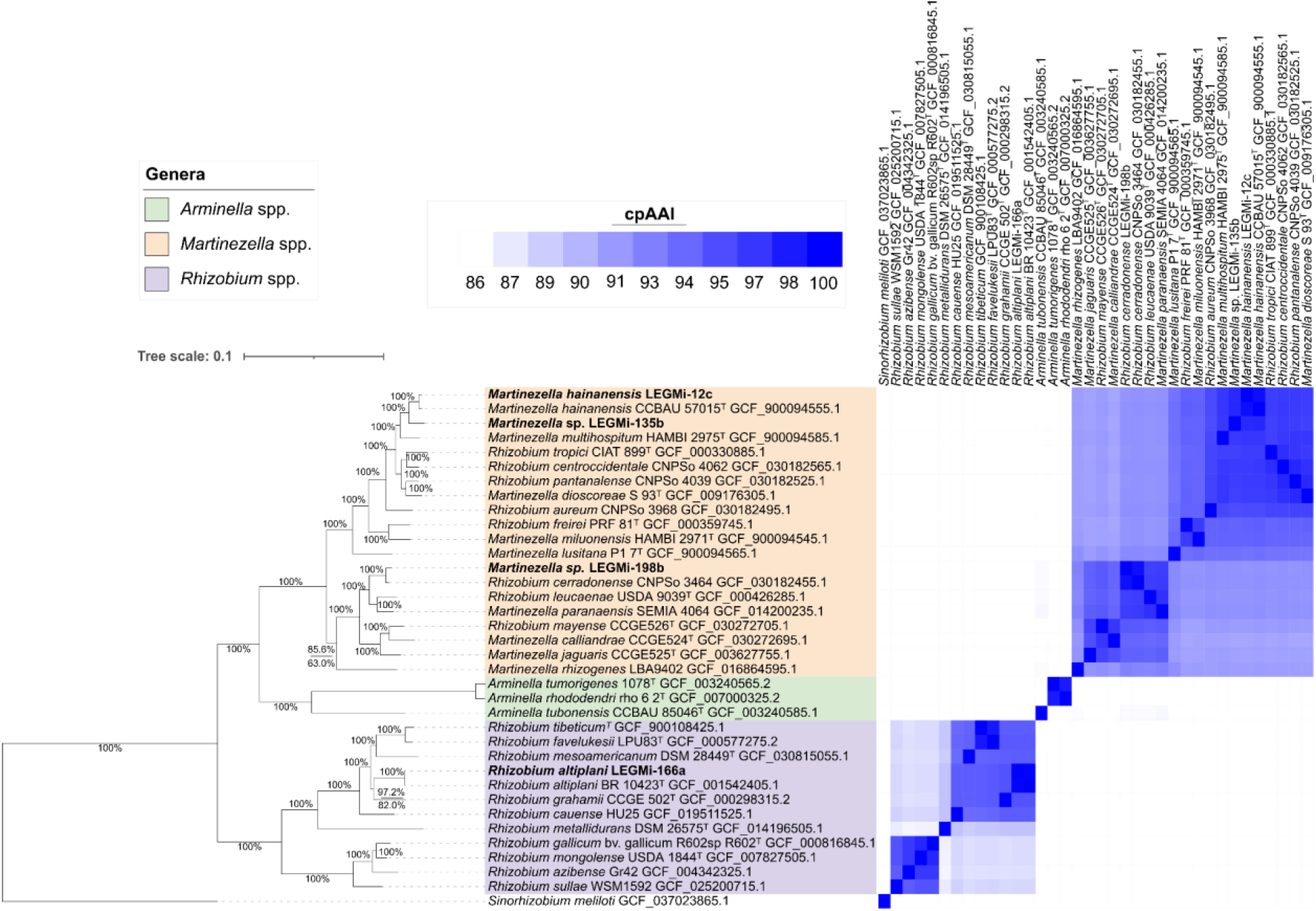
Maximum-likelihood phylogenomic tree inferred from 170 conserved marker proteins of the genera *Arminella*, *Martinezella*, and *Rhizobium* (left), and core proteome average amino acid identity (cpAAI) matrix calculated from the same set of 170 conserved proteins (right). The phylogenomic tree was reconstructed using IQ-TREE v2.2.0 under the LG+F+R5 substitution model. A total of 36 genomes representing the genera *Rhizobium*, *Martinezella*, and *Arminella*, corresponding to the closest relatives of the isolates characterized in this study, were included in the analysis. Genome accession numbers are provided next to each strain name, and type strains are indicated by the superscript “T”. Branch support values correspond to SH-aLRT and ultrafast jackknife support. When both methods recovered a support value of 100, a single value (100) is shown; otherwise, both support values are displayed in the format SH-aLRT/ultrafast jackknife. The scale bar represents the average number of amino acid substitutions per site. The three putative *Martinezella* strains and the *Rhizobium* species identified in this study are shown in bold. The tree was rooted using *Sinorhizobium meliloti* as the outgroup. In the cpAAI matrix, values below the genus delimitation threshold (86.5%) are shown in white, whereas values equal to or greater than 86.5% are represented using a gradient of blue shades, with darker colors indicating higher cpAAI values up to 100%. Tree visualization and annotation were performed using iTOL v6.

Based on their cpAAI values, which exceed the 86.5% threshold for genus delineation, and their position in phylogenomic trees within the *Martinezella* clade, we suggest that it would be appropriate to reclassify the following species to the genus *Martinezella*: *Rhizobium leucaenae* USDA 9039^T^ (GCF_000426285.1), *R. tropici* CIAT 899^T^ (GCF_000330885.1), *R. centroccidentale* CNPSo 4062 (GCF_030182565.1), *R. pantanalense* CNPSo 4039 (GCF_030182525_1), *R. aureum* CNPSo 3968 (GCF_030182495.1), *R. freirei* PRF 81^T^ (GCF_000359745.1), and *R. mayense* CCGE526T (GCF_030272705.1). The cpAAI phylogenetic tree files and the concatenated protein sequences used for the analysis are provided as supplementary material.

The ANI and dDDH values for strain LEGMi-135b (ANI: 94%, dDDH: 53.2%) relative to *M. hainanensis* CCBAU-57015T, values below the commonly accepted species thresholds (ANI ≤ 95–96% and dDDH ≤ 70%), suggest that strain LEGMi-135b represents a potentially new species within the genus *Martinezella* (Tables 2 and 3). Strain LEGMi-12c is related to the species *M. hainanensis* CCBAU-57015^T^ (ANI: 98%, dDDH: 87%), LEGMi-198b to *R. cerradonense* CNPSo 3464 (ANI: 98%, dDDH: 81.4%), and LEGMi-166a to *R. altiplani* BR 10423^T^ (ANI: 100%, dDDH: 99.3%).

**Table 3.** dDDH (d4) values (in %) between *Martinezella* and *Rhizobium* strains under study and their closest published species.

|  | *Martinezella hainanensis*<br>CCBAU- 57015<sup>T</sup>-<br>GCF_900094555.1 | *Rhizobium altiplani*<br>BR 10423<sup>T</sup>-<br>GCF_001542405.1 | *Rhizobium*<br>*cerradonense* CNPSO<br>3464- GCF_030182455.1 |
| --- | --- | --- | --- |
| *Martinezella* sp. LEGMi-12c | 87.00 | 21.20 | 25.60 |
| *Martinezella* sp. LEGMi-135b | 53.20 | 21.50 | 25.50 |
| *Rhizobium* sp. LEGMi-166a | 21.10 | 99.30 | 21.60 |
| *Martinezella* sp. LEGMi-198b | 25.60 | 21.70 | 81.40 |

**Table 4.** ANI values (in %) between *Martinezella* and *Rhizobium* strains under study and their closest published species.

|  | *Martinezella hainanensis*<br>CCBAU- 57015<sup>T</sup>-<br>GCF_900094555.1 | *Rhizobium altiplani*<br>BR 10423<sup>T</sup>-<br>GCF_001542405.1 | *Rhizobium*<br>*cerradonense* CNPSO<br>3464- GCF_030182455.1 |
| --- | --- | --- | --- |
| *Martinezella* sp. LEGMi-12c | 98 | 84 | 86 |
| *Martinezella* sp. LEGMi-135b | 94 | 84 | 86 |
| *Rhizobium* sp. LEGMi-166a | 84 | 100 | 85 |
| *Martinezella* sp. LEGMi-198b | 86 | 84 | 98 |

A phylogenetic tree based on the maximum likelihood method using 16S rRNA sequences shows strain LEGMi-12c clustering near *M. multihospitum* HAMBI 2975^T^ and *R. frieri* PRF 81^T^, while LEGMi-135b forms a separate but closely related branch within the *Martinezella* clade (Fig. 4). Strain LEGMi-198b is also found within the *Martinezella* clade, closely related to *R. cerradonense* CNPSo 3464. In contrast, LEGMi-166a and *R. altiplani* BR 10423^T^ form a sister clade to *R. mesoamericanum* STM3625, yet fall within the “leguminosarum-etli” clade.

**Fig. 4.**
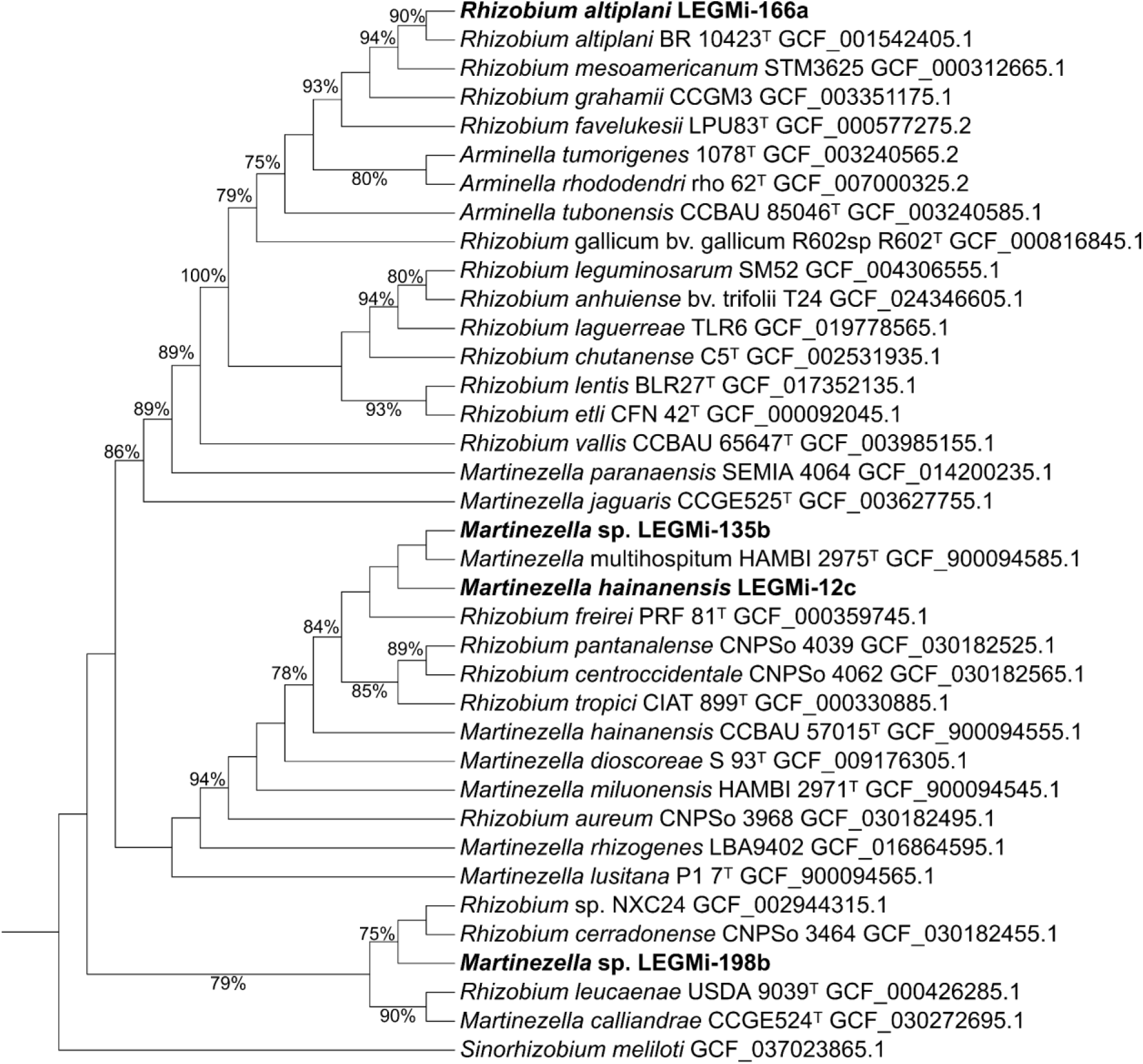
Maximum-likelihood phylogenetic tree based on the *16S rRNA* gene. The phylogeny was inferred using IQ-TREE v2.2.0 under the TIM3+F+I+G4 substitution model. The analysis included 37 genomes belonging to the genera *Rhizobium*, *Martinezella*, and *Arminella*, representing the closest relatives of the isolates examined in this study. Genome accession numbers are indicated alongside each strain name. Type strains are denoted by the superscript “T”, and bootstrap support values ≥60% are shown in the corresponding nodes.The three putative *Martinezella* strains and the *Rhizobium* species identified in this study are highlighted in bold. The genome of *Sinorhizobium meliloti* was used as the outgroup to root the tree. Branch lengths were omitted to improve tree visualization. Tree visualization and annotation were performed using iTOL v6.

BLASTP analysis of candidate proteins recovered from the genomes of strains LEGMi-135b and LEGMi-12c identified proteins annotated as GT2-family glycosyltransferases as the best matches, with no proteins annotated as *nodC* being recovered. Consequently, these sequences were excluded from the phylogenetic analysis of *nodC* genes extracted from the whole genomes. This aligns with results from functional annotation tools such as eggnog-mapper and RAST, which did not functionally annotate the *nodC* gene in these genomes, although they did identify *in silico* genes related to nodulation, nodulation regulation, and nitrogen fixation. A search for orthologs using *nodC* reference sequences from UniProt allowed for more specific gene identification; thus, we conclude that the genome either lacks the *nodC* gene, lost it, or that the gene was lost during the assembly process (given that the data came from a draft assembly). Contradictorily, the nodulation assay with *P. vulgaris* cv. Cabécar yielded positive results for strain LEGMi-12c (eleven nodules) in one of the two test replicates, but not for LEGMi-135b (zero nodules), across the four test replicates (Table 5).

**Table 5.**
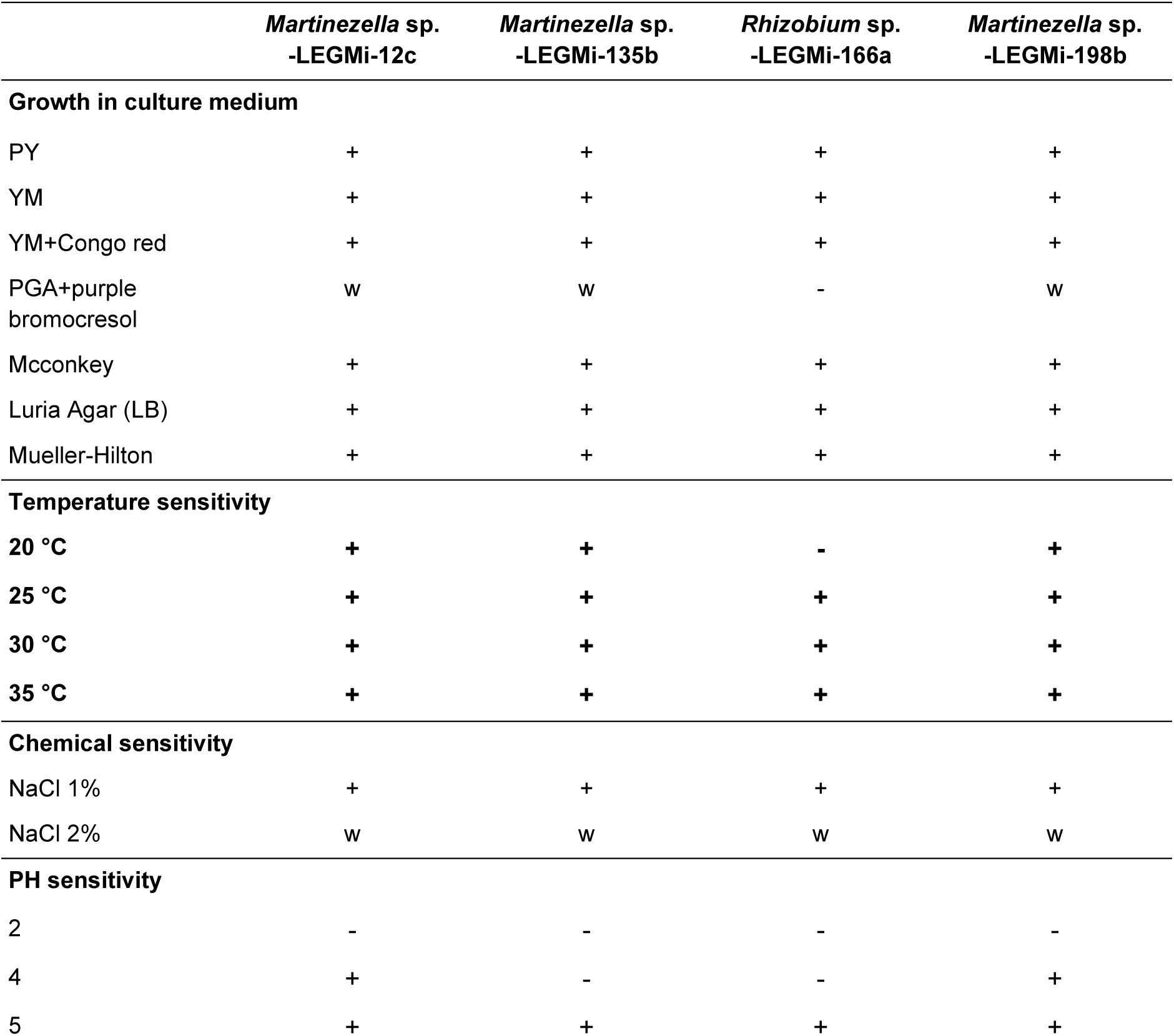
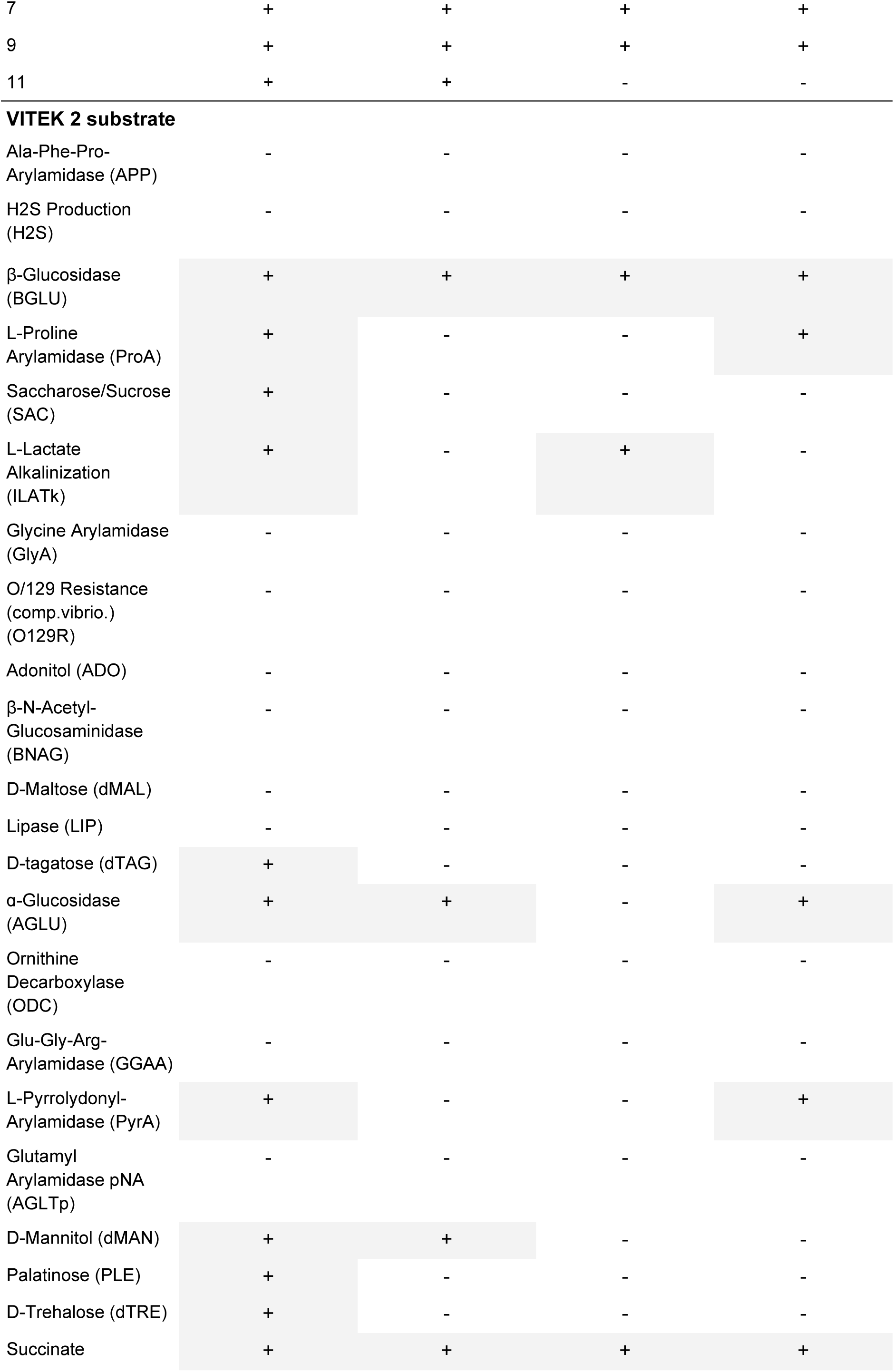
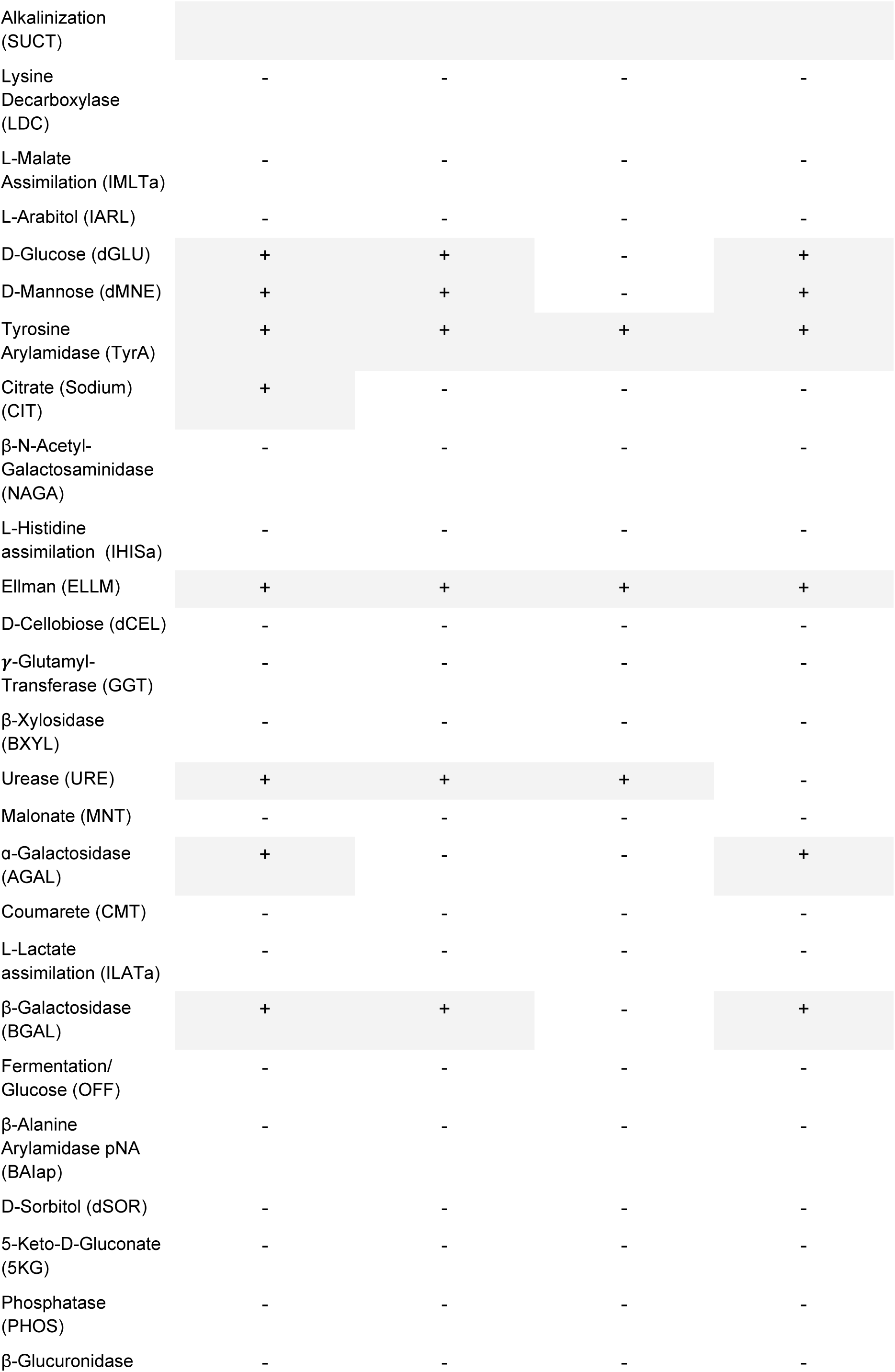
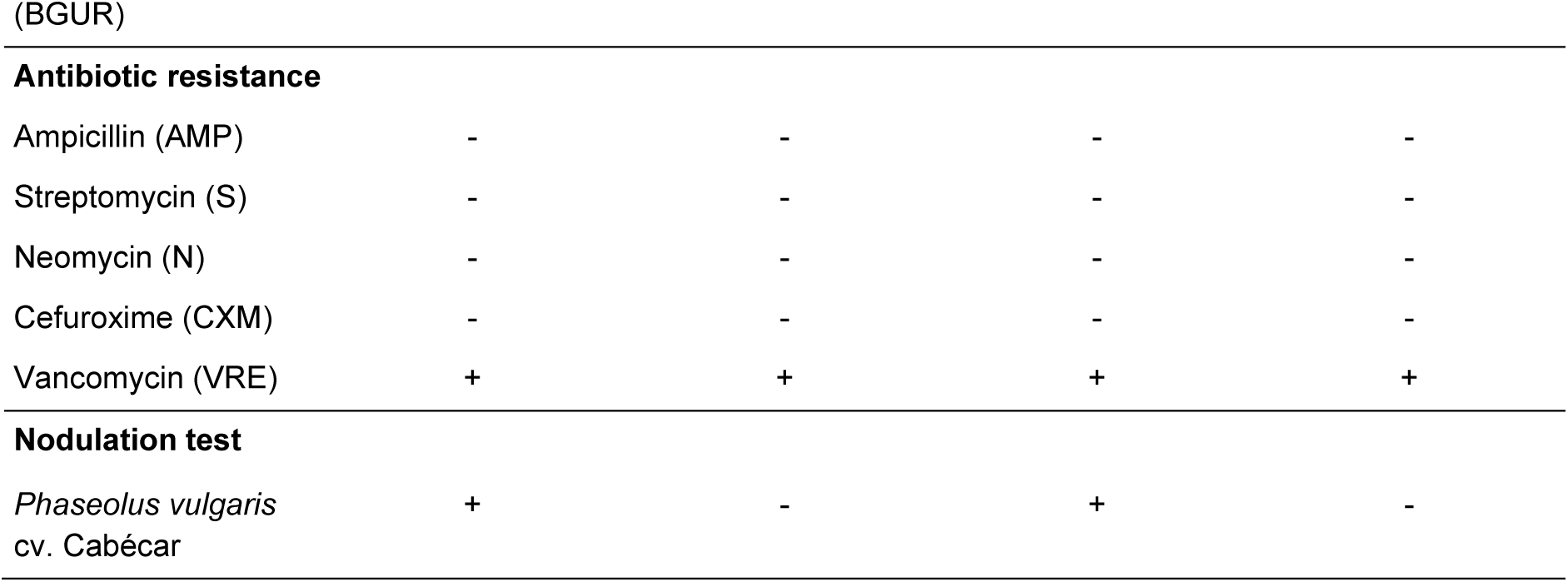
Phenotypic features of the isolated strains. −, Negative; +, positive; w, weak reaction.

|  | *Martinezella* sp.<br>-LEGMI-12c | *Martinezella* sp.<br>-LEGMI-135b | *Rhizobium* sp.<br>-LEGMI-166a | *Martinezella* sp.<br>-LEGMI-198b |
| --- | --- | --- | --- | --- |
| **Growth in culture medium** |  |  |  |  |
| PY | + | + | + | + |
| YM | + | + | + | + |
| YM+Congo red | + | + | + | + |
| PGA+purple<br>bromocresol | w | w | - | w |
| Mcconkey | + | + | + | + |
| Luria Agar (LB) | + | + | + | + |
| Mueller-Hilton | + | + | + | + |
| **Temperature sensitivity** |  |  |  |  |
| 20 °C | + | + | - | + |
| 25 °C | + | + | + | + |
| 30 °C | + | + | + | + |
| 35 °C | + | + | + | + |
| **Chemical sensitivity** |  |  |  |  |
| NaCl 1% | + | + | + | + |
| NaCl 2% | w | w | w | w |
| **PH sensitivity** |  |  |  |  |
| 2 | - | - | - | - |
| 4 | + | - | - | + |
| 5 | + | + | + | + |
| 7 | + | + | + | + |
| 9 | + | + | + | + |
| 11 | + | + | - | - |
| **VITEK 2 substrate** |  |  |  |  |
| Ala-Phe-Pro-Arylamidase (APP) | - | - | - | - |
| H<sub>2</sub>S Production (H<sub>2</sub>S) | - | - | - | - |
| β-Glucosidase (BGLU) | + | + | + | + |
| L-Proline Arylamidase (ProA) | + | - | - | + |
| Saccharose/Sucrose (SAC) | + | - | - | - |
| L-Lactate Alkalinization (ILATk) | + | - | + | - |
| Glycine Arylamidase (GlyA) | - | - | - | - |
| O/129 Resistance (comp.vibrio.) (O129R) | - | - | - | - |
| Adonitol (ADO) | - | - | - | - |
| β-N-Acetyl-Glucosaminidase (BNAG) | - | - | - | - |
| D-Maltose (dMAL) | - | - | - | - |
| Lipase (LIP) | - | - | - | - |
| D-tagatose (dTAG) | + | - | - | - |
| α-Glucosidase (AGLU) | + | + | - | + |
| Ornithine Decarboxylase (ODC) | - | - | - | - |
| Glu-Gly-Arg-Arylamidase (GGAA) | - | - | - | - |
| L-Pyrrolydonyl-Arylamidase (PyrA) | + | - | - | + |
| Glutamyl Arylamidase pNA (AGLTp) | - | - | - | - |
| D-Mannitol (dMAN) | + | + | - | - |
| Palatinose (PLE) | + | - | - | - |
| D-Trehalose (dTRE) | + | - | - | - |
| Succinate | + | + | + | + |
| Alkalinization<br>(SUCT) |  |  |  |  |
| Lysine<br>Decarboxylase<br>(LDC) | - | - | - | - |
| L-Malate<br>Assimilation (IMLTa) | - | - | - | - |
| L-Arabitol (IARL) | - | - | - | - |
| D-Glucose (dGLU) | + | + | - | + |
| D-Mannose (dMNE) | + | + | - | + |
| Tyrosine<br>Arylamidase (TyrA) | + | + | + | + |
| Citrate (Sodium)<br>(CIT) | + | - | - | - |
| β-N-Acetyl-Galactosaminidase<br>(NAGA) | - | - | - | - |
| L-Histidine<br>assimilation (IHISa) | - | - | - | - |
| Ellman (ELLM) | + | + | + | + |
| D-Cellobiose (dCEL) | - | - | - | - |
| γ-Glutamyl-Transferase (GGT) | - | - | - | - |
| β-Xylosidase<br>(BXYL) | - | - | - | - |
| Urease (URE) | + | + | + | - |
| Malonate (MNT) | - | - | - | - |
| α-Galactosidase<br>(AGAL) | + | - | - | + |
| Coumarete (CMT) | - | - | - | - |
| L-Lactate<br>assimilation (ILATa) | - | - | - | - |
| β-Galactosidase<br>(BGAL) | + | + | - | + |
| Fermentation/<br>Glucose (OFF) | - | - | - | - |
| β-Alanine<br>Arylamidase pNA<br>(BAIap) | - | - | - | - |
| D-Sorbitol (dSOR) | - | - | - | - |
| 5-Keto-D-Gluconate<br>(5KG) | - | - | - | - |
| Phosphatase<br>(PHOS) | - | - | - | - |
| β-Glucuronidase | - | - | - | - |
| (BGUR) |  |  |  |  |
| **Antibiotic resistance** |  |  |  |  |
| Ampicillin (AMP) | - | - | - | - |
| Streptomycin (S) | - | - | - | - |
| Neomycin (N) | - | - | - | - |
| Cefuroxime (CXM) | - | - | - | - |
| Vancomycin (VRE) | + | + | + | + |
| **Nodulation test** |  |  |  |  |
| *Phaseolus vulgaris*<br>cv. Cabécar | + | - | + | - |

Regarding the *nodC* gene of LEGMi-198b, it belongs to the clade containing *R. tropici* CIAT 899^T^ and *R. cerradonense* CNPSo 3464, yet it lies on a branch independent from them (Fig. 5, Fig. S3). However, no nodulation by this strain was observed in the four replicates of the assay with *P. vulgaris* cv. Cabécar plants, despite the *nodC* gene having been detected *in silico* during functional annotation (Table 5). Moura et al. (49) isolated the type strain from common bean (*P. vulgaris*) nodules and demonstrated that it was capable of nodulating its original host as well as *Macroptilium atropurpureum*. Differences in this capability could be attributed to variations in the cultivar used in our test. Strain LEGMi-166a is phylogenetically related to *R. altiplani* BR 10423^T^. When evaluating the nodulation capacity of strain LEGMi-166a on *P. vulgaris* cv. Cabécar plants, positive nodulation was observed, with abundant nodule formation (87–96 nodules per plant) in both test replicates. Both functional annotation and the analysis of putative genes enabled the *in silico* identification of the *nodC* gene within the assembled genome. Baraúna et al. (50) reported that the type strain (*R. altiplani* BR 10423^T^) is capable of nodulating *Mimosa pudica* (its original isolation host), *Crotalaria juncea*, *Vigna unguiculata*, and *P. vulgari*s.

**Fig. 5.**
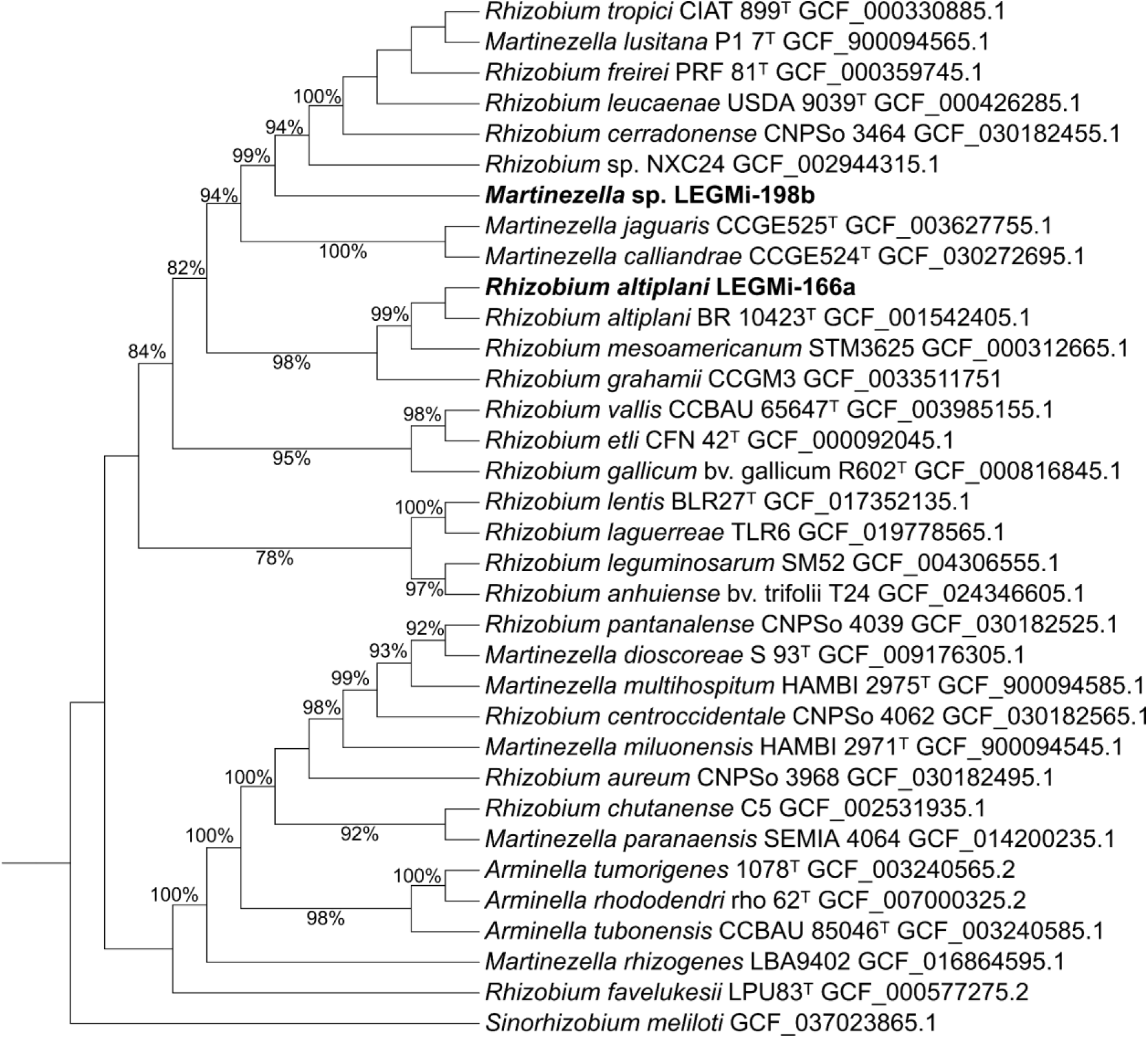
Maximum-likelihood phylogenetic tree inferred from the *nodC* gene. The phylogeny was reconstructed using IQ-TREE v2.2.0 under the Q.pfam+F+R4 substitution model. The analysis included 34 genomes belonging to the genera *Rhizobium*, *Martinezella*, and *Arminella*, representing the closest relatives of the isolates examined in this study. Genome accession numbers are indicated next to each strain name. Type strains are denoted by the superscript “T”, and bootstrap support values ≥70% are shown in the corresponding nodes.The three putative *Martinezella* and the *Rhizobium* species identified in this study are highlighted in bold.Branch lengths were omitted to improve tree visualization. The genome of *Sinorhizobium meliloti* was used as the outgroup to root the tree. Tree visualization and annotation were performed using iTOL v6.

### Physiology and chemotaxonomy

The four strains in the study were able to grow on YM, PY, MacConkey, LB, and Mueller-Hinton media (Table 5). On YMA supplemented with Congo red, all four strains exhibited growth, forming mucoid colonies with a translucent pinkish tone. In contrast, on PGA medium supplemented with bromocresol purple, strains LEGMI-12c, LEGMI-135b, and LEGMI-198b showed weak, slow growth, while LEGMI-166a showed no growth at all. On YMA medium supplemented with bromothymol blue, all four strains were able to grow and acidify the medium, a change observed as a shift to yellow.

All strains in the study are rod-shaped, aerobic, Gram-negative bacteria, approximately 3 μm in size (Fig. S4). When cultured on YMA, the colonies appear translucent creamy-white and convex, with a mucilaginous texture after 48 hours. Strains LEGMI-12c, LEGMI-135b, and LEGMI-198b were able to grow within the temperature range of 20°C to 35°C; however, LEGMI-166a grew only between 25°C and 35°C. All four strains grew on YMA supplemented with 1% NaCl, and showed weak growth on YMA with 2% NaCl (Table 5). LEGMI-12c was able to grow at pH 4– 11, LEGMI-135 at pH 5–11, LEGMI-166a at pH 5–9, and LEGMI-198b at pH 4–9 (Table 5, Fig. S5). None of the strains were able to grow in YM medium at pH 2. All strains were sensitive to the antibiotics ampicillin (AMP), streptomycin (S), neomycin (N), and cefuroxime (CXM), but were able to grow in medium supplemented with vancomycin (Table 5).

The four strains under study shared Ellman (ELLM), Tyrosine Arylamidase (TyrA), Succinate Alkalinization (SUCT), and β-Glucosidase (BGLU) enzymatic activities. Meanwhile, the three strains identified as belonging to the genus *Martinezella* shared the ability to utilize D-Glucose (dGLU) and D-Mannose (dMNE) as carbon sources and exhibited β-Galactosidase (BGAL), β-Glucosidase (BGLU), and α-Glucosidase (AGLU) activities.

Based on the genetic and phenotypic data presented in this study, we propose that strain LEGMi-135b represents a new species named *Martinezella legmii* sp. nov. Likewise, strain LEGMi-198b represents a new species of the genus *Martinezella*, named *Martinezella ivanii* sp. nov.

### Description of *Martinezella legmii* sp. nov

*Martinezella legmii* (leg’mi.i. N.L. gen. n. *legmii*, of LEGMi, named after the LEGMi Laboratory, where the strain was isolated and characterized).

Fast growing, aerobic, non spore forming, rodshaped, Gram-negative bacteria approximately 3 μm in size. When cultured on YMA, the colonies appear translucent creamy white and convex, with a mucilaginous texture after 48 hours. Optimal growth occurs at pH 7 and 28°C, though it can grow within a temperature range of 25–35°C and a pH range of 4–11. Strain LEGMi-135b grows on YMA with 1% NaCl but exhibits slow growth on YMA with 2% NaCl. The strain is capable of utilizing carbon sources such as D-mannitol (dMAN), D-glucose (dGLU), and D-mannose (dMNE), but not sucrose (SAC), adonitol (ADO), D-maltose (dMAL), D-tagatose (dTAG), palatinose (PLE), D-trehalose (dTRE), L-arabitol (IARL), D-cellobiose (dCEL), D-sorbitol (dSOR), or 5-keto-D-gluconate (5KG). It showed no positive response to organic acids such as L-lactate alkalinization (ILATk), L-malate assimilation (IMLTa), citrate (sodium) (CIT), malonate (MNT), coumarate (CMT), or L-lactate assimilation (ILATa). It exhibits positive enzymatic activity for β-glucosidase (BGLU), α-glucosidase (AGLU), succinate alkalinization (SUCT), tyrosine arylamidase (TyrA), Ellman (ELLM), urease (URE), and β-galactosidase (BGAL). It is susceptible to the antibiotics kanamycin, streptomycin, neomycin, and cefuroxime, but shows resistance to vancomycin.

The type strain is LEGMi-135b^T^; it was isolated from nodules of *Lonchocarpus felipei* grown in a nursery in San Carlos, Heredia, Costa Rica. *Martinezella legmii* LEGMi-135b^T^ does not show nodulation ability in *in vivo* experiments on *Phaseolus vulgaris* cv. Cabécar seedlings. *Martinezella legmii* LEGMi-135b^T^ has a genome size of 6,62 Mb, with 6,192 predicted genes and a DNA G+C content of 59.7 mol%.

### Description of *Martinezella ivanii* sp. nov

*Martinezella ivanii* (i.va’ni.i. N.L. gen. masc. n. ivanii, of Ivan, named in memory of Ivan, the late brother of the author).

Fast growing, aerobic, non spore forming, rodshaped, Gram-negative bacteria approximately 3 μm in size. When cultured on YMA, the colonies appear translucent creamy white and convex with a mucilaginous texture after 48 hours. Optimal growth occurs at pH 7 and 28°C; it can grow within a temperature range of 25°C to 35°C and a pH range of 5 to 9. It grows on YMA with 1% NaCl but shows weak growth on YMA with 2% NaCl. The strain can utilize carbon sources such as D-Glucose (dGLU) and D-Mannose (dMNE), but not Sucrose (SAC), Adonitol (ADO), D-Maltose (dMAL), D-Tagatose (dTAG), D-Mannitol (dMAN), Palatinose (PLE), D-Trehalose (dTRE), L-Arabitol (IARL), D-Cellobiose (dCEL), D-Sorbitol (dSOR), or 5-Keto-D-Gluconate (5KG). It showed no positive response to organic acids such as L-Lactate Alkalinization (ILATk), L-Malate Assimilation (IMLTa), Citrate (CIT), Malonate (MNT), Coumarate (CMT), L-Lactate Assimilation (ILATa), or L-Histidine Assimilation (IHISa). It exhibits positive enzymatic activity for β-Glucosidase (BGLU), α-Glucosidase (AGLU), Succinate Alkalinization (SUCT), Tyrosine Arylamidase (TyrA), Ellman (ELLM), L-Pyrrolydonyl-Arylamidase (PyrA), α-Galactosidase (AGAL), and β-Galactosidase (BGAL). It is susceptible to the antibiotics Kanamycin, Streptomycin, Neomycin, and Cefuroxime, and is resistant to Vancomycin.

The type strain is LEGMi-198b^T^; it was isolated from nodules of *Calliandra haematocephala* grown in a nursery in Heredia, Costa Rica. *Martinezella ivanii* LEGMi-198b^T^ does not show nodulation ability in *in vivo* experiments on *Phaseolus vulgaris* cv. Cabécar seedlings, although the *nodC* gene is detected *in silico*. *Martinezella ivanii* LEGMi-198b has a genome size of 6,83 Mb, with 6,357 predicted genes and a DNA G+C content of 59.53 mol%.

## Abbreviations

ANI: Average nucleotide identity
cpAAI: Core-protein Amino Acid Identity
dDDH: digital DNA–DNA hybridization
GTDB: Genome Taxonomy Database
LEGMi: Laboratory of Genetics and Ecology of Micro-organisms
NCBI: National Center for Biotechnology Information
PGAP: NCBI Prokaryotic Genome Annotation Pipeline
SCCGs: shared core single-copy genes
SRA: Sequence Read Archive
TEM: Transmission electron microscopy
TYGS: Type Strain Genome Server.

## Funding information

This study was funded by the University of Costa Rica (projects B9-204 and C0-524) and by the Ministerio de Ciencia, Innovación, Tecnología y Telecomunicaciones de Costa Rica (MICITT)-Consejo Nacional para Investigaciones Científicas y Tecnológicas (CONICIT) (project FI-036B-19).

## Acknowledgment

We acknowledge the Colaboratorio Nacional de Computación Avanzada at the Centro Nacional de Alta Tecnología (CeNAT), Costa Rica, for providing access to the Kabré supercomputer facilities. We also thank the Faculty of Microbiology at the University of Costa Rica, particularly PhD. Pamela Altamirano Silva and Yeimmy Ramírez Zúñiga, for performing the VITEK 2 analyses. We further acknowledge the Laboratory of Pharmaceutical Analysis and Consulting of the Institute for Pharmaceutical Research (INIFAR), Faculty of Pharmacy, University of Costa Rica, particularly Jeison Corrales Sánchez and Arlene Loria Gutiérrez, for carrying out the lyophilization of the strains.

## Author contributions

Conceptualization, D.C.P., K.R.J. and B.M.G; methodology, D.C.P., E.A.S., K.R.J. and B.M.G; formal analysis, D.C.P.; investigation, D.C.P.; writing – original draft preparation, D.C.P.; supervision, E.A.S., K.R.J., M.V.A. and B.M.G.; visualization, D.C.P. and I.R.R.; funding acquisition, K.R.J.; writing, review and editing, D.C.P., E.A.S., K.R.J. and B.M.G. All authors have read and agreed to the published version of the manuscript.

## Conflicts of interest

The authors declare that there are no conflicts of interest.

## Notes

### Competing Interest Statement

The authors have declared no competing interest.

