## supplementary figs and tables for "Characterization of two *Martinezella* spp. species: *M. legmii* sp. nov. and *M. ivanii* sp. nov., from tropical legume trees in Costa Rica"

### Supplemental material

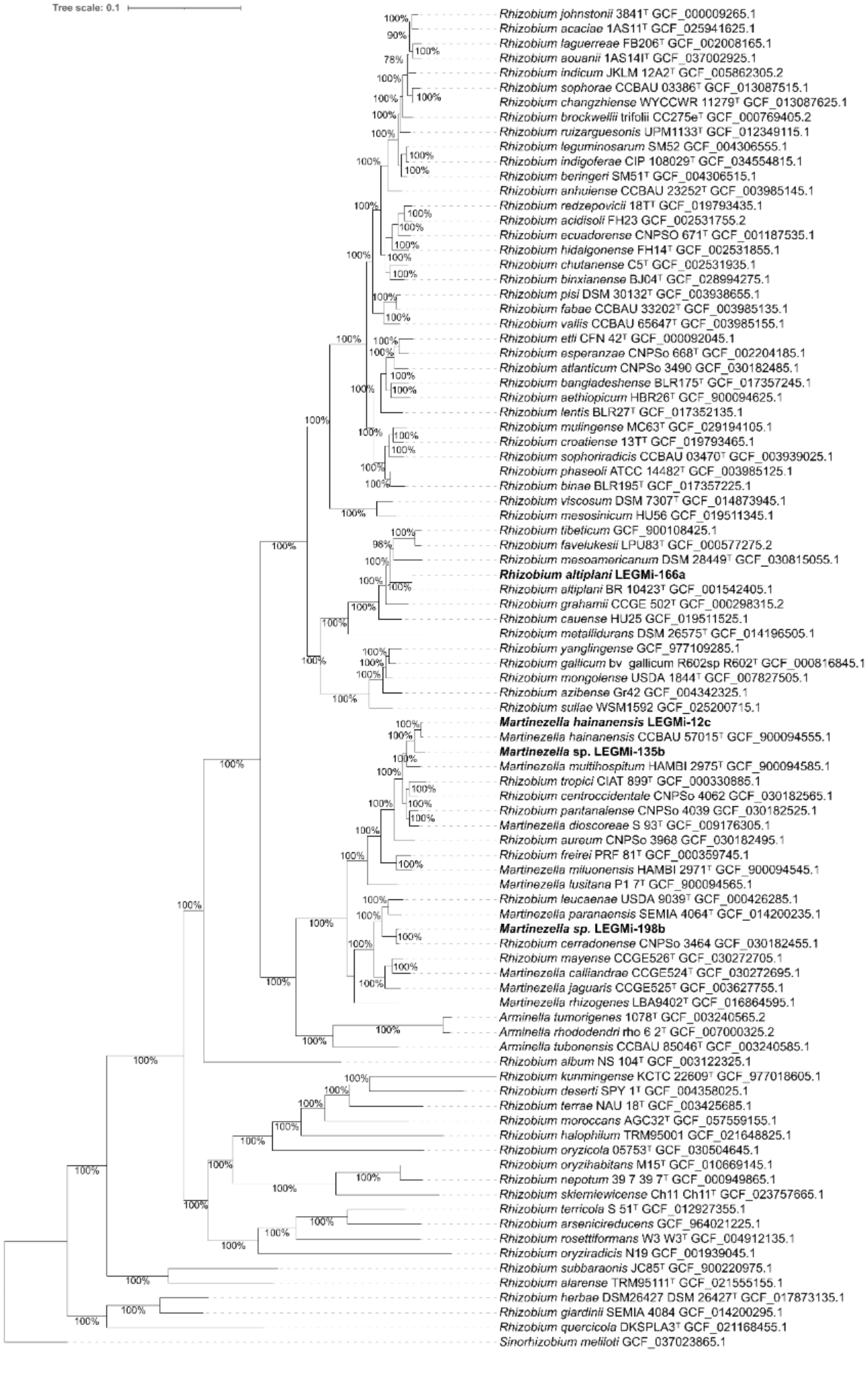

Figure S1. Phylogenomic tree inferred from the core genome of 91 bacterial genomes, comprising 651 single-copy orthologous gene sequences and a total of 59,241 genes. Phylogenetic reconstruction was performed using FastTree under the JTT-CAT substitution model with 1,000 bootstrap replicates. Reference genomes were manually selected from the GenBank database, prioritizing type strains and including the closest available genomes of *Rhizobium*, *Martinezella* and *Arminella*, identified via the List of Prokaryotic names with Standing in Nomenclature (LPSN), as well as genomes of species that, according to the LPSN, have a "not validly published" nomenclatural status but serve as reference genomes not associated with type material.Genome accession numbers are indicated next to the name of each strain. Type strains are marked with a superscript "T", while bootstrap support values ≥70% are shown at the corresponding nodes. The three putative *Martinezella* and the *Rhizobium* species identified in this study are highlighted in bold. The genome of *Sinorhizobium meliloti* was used to root the tree. Tree visualization and annotation were carried out using iTOL v6.

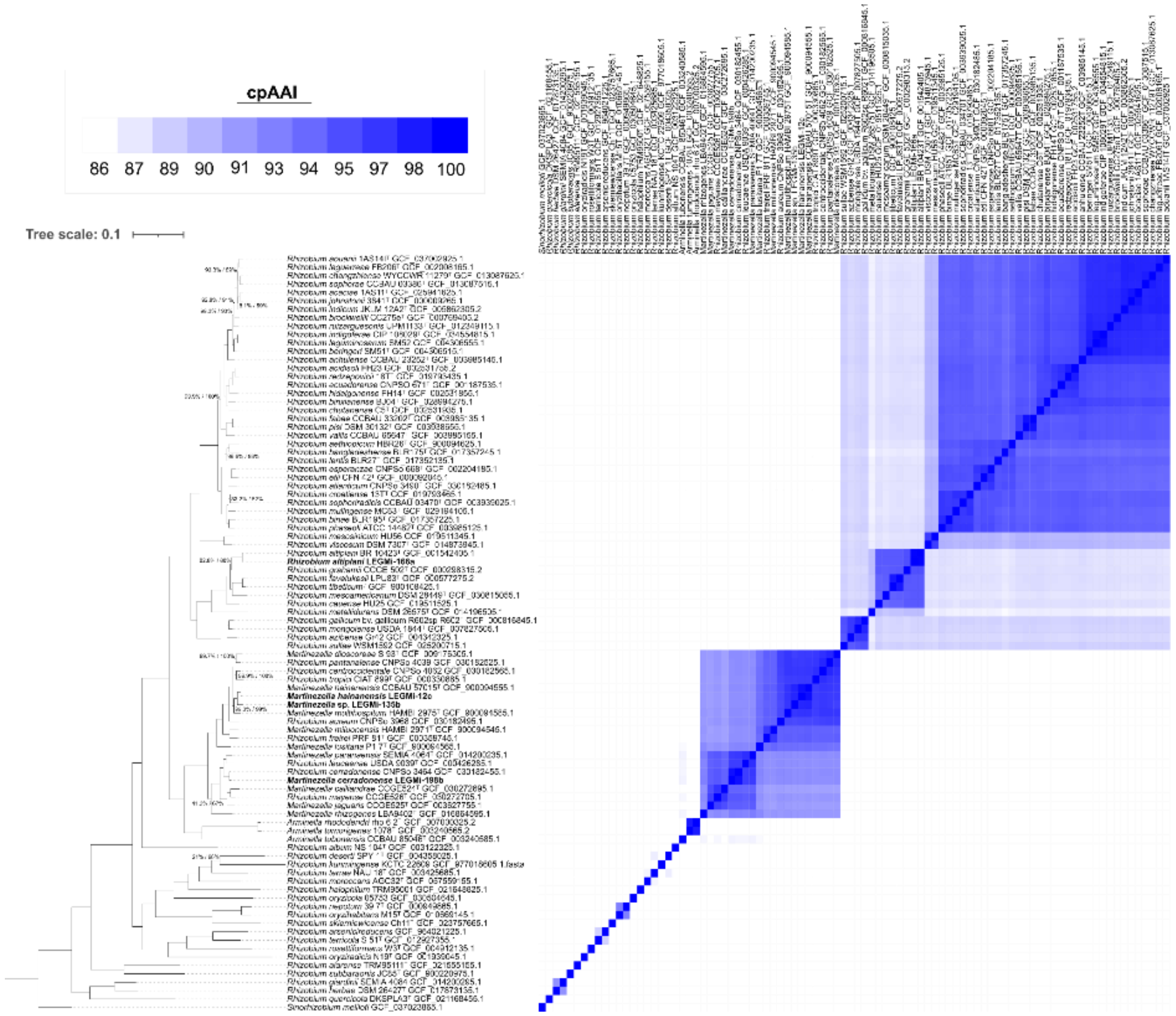


Figure S2. Maximum-likelihood phylogenomic tree inferred from 170 conserved marker proteins of the genera *Arminella*, *Martinezella*, and *Rhizobium* (left), and core proteome average amino acid identity (cpAAI) matrix calculated from the same set of

170 conserved proteins (right). The phylogenomic tree was reconstructed using IQ-TREE v2.2.0 under the LG+F+R5 substitution model. The analysis included a total of 86 genomes representing the genera *Rhizobium*, *Martinezella*, and *Arminella*. All available reference genomes for these genera were considered, with preference given to type strains when available. Genome accession numbers are provided next to each strain name, and type strains are indicated by the superscript "T". Branch support values correspond to SH-aLRT and ultrafast jackknife support. When both methods recover a support value of 100, a single value (100) is shown; otherwise, both support values are displayed in the format SH-aLRT/ultrafast jackknife. The scale bar represents the average number of amino acid substitutions per site. The three putative *Martinezella* strains and the *Rhizobium* species identified in this study are shown in bold. The tree was rooted using *Sinorhizobium meliloti* as the outgroup. In the cpAAI matrix, values below the genus delimitation threshold (86.5%) are shown in white, whereas values equal to or greater than 86.5% are represented using a gradient of blue shades, with darker colors indicating higher cpAAI values up to 100%. Tree visualization and annotation, including the cpAAI matrix, were performed using iTOL v6.

Table S1. Identification of nodulation and nitrogen fixation related genes in the genomes of *Martinezella* sp. LEGMI-12c, *Martinezella* sp. LEGMI-135B, *Rhizobium* sp. LEGMI-166a, and *Martinezella* sp. LEGMI-198b based on GenBank genome annotations generated using the NCBI Prokaryotic Genome Annotation Pipeline (PGAP). The table shows the presence or absence of the *nodA*, *nodB*, *nodC*, *fixH*, and *nifH* genes. Gene symbols and corresponding locus tags are provided for genes identified in each genome. “--” indicates that no corresponding gene was identified in the GenBank PGAP annotation.

| **Symbol** | ***Martinezella* sp. LEGMI-12c** | ***Martinezella* sp. LEGMI-135B** | ***Rhizobium* sp. LEGMI-166a** | ***Martinezella* sp. LEGMI-198b** |
|---|---|---|---|---|
| *nodA* | -- | -- | -- | ACXHXG_RS19385 |
| *nodB* | -- | -- | -- | ACXHXG_RS19390 |
| *nodC* | -- | -- | -- | ACXHXG_RS19395 |
| *fixH* | ACXHMN_RS12015 | -- | ACXHXM_RS34975 | ACXHXG_RS32050 |
| *nifH* | -- | -- | ACXHXM_RS33670 | ACXHXG_RS19590 |

Table S2. GTDB-Tk genome-based taxonomic classification of the analyzed *Martinezella* and *Rhizobium* strains.

| **Strain** | **Classification** | **FastANI Reference** | **FastANI Reference Radius** |
|---|---|---|---|
| LEGMi-166a | *Rhizobium altiplani* | GCF_001542405.1 | 95 |
| LEGMi-198b | *Rhizobium* sp002944315 | GCF_002944315.1 | 95 |
| LEGMi-12c | *Martinezella hainanense* | GCF_900094555.1 | 95 |
| LEGMi-135b | *Martinezella* sp. | - | - |

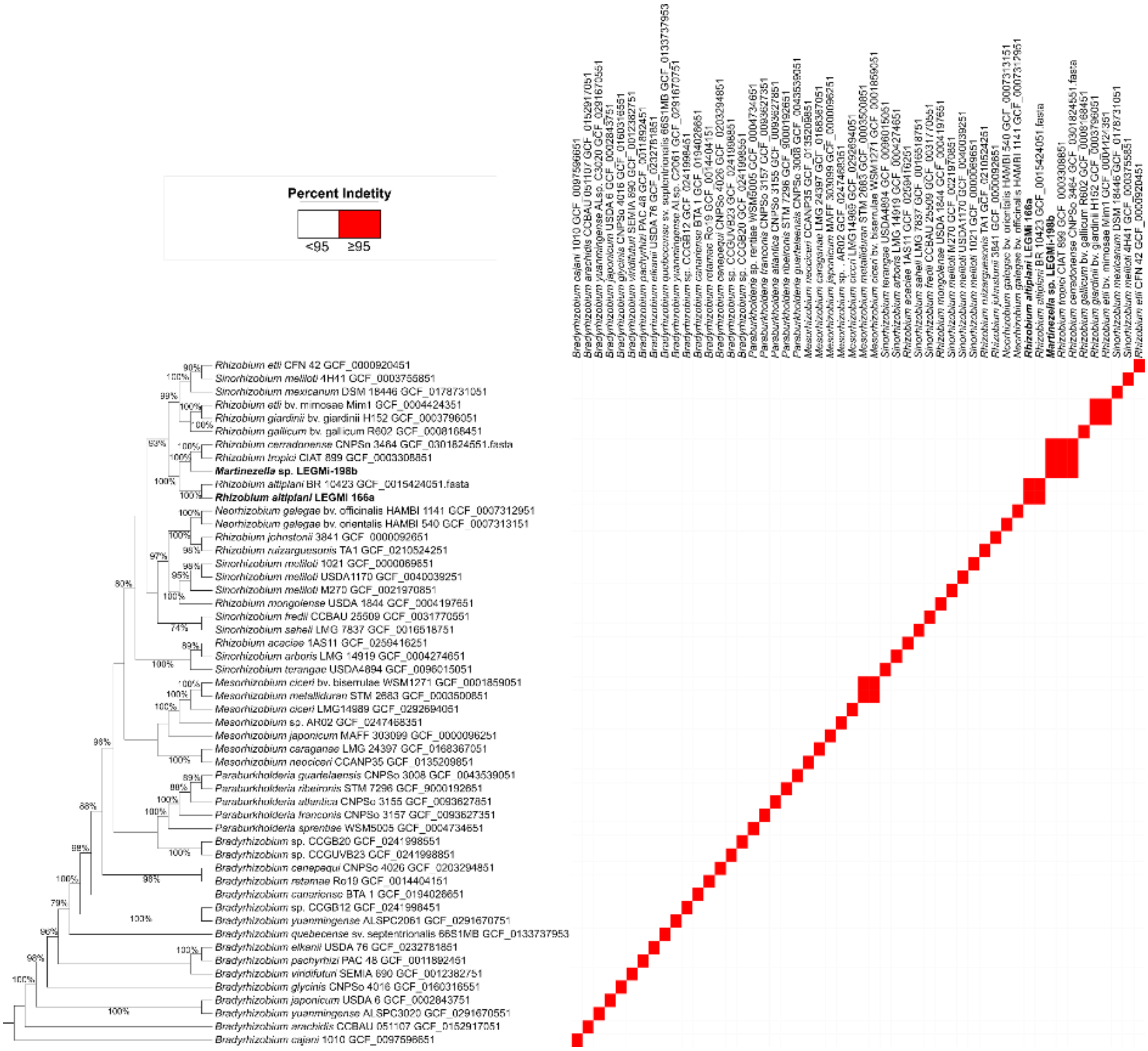

Figure S3. Maximum-likelihood phylogenetic tree inferred from the *nodC* gene. The phylogeny was reconstructed using IQ-TREE v2.2.0 under the Q.mammal+I+G4 substitution model. The analysis included 50 genomes belonging to the reference genomes of the symbiovars compiled by Martínez-Romero et al. (2024), along with the strains LEGMi-166a and LEGMi-198b. Genome accession numbers are indicated next to each strain name. Type strains are denoted by the superscript "T", and bootstrap support values ≥70% are shown at the corresponding nodes. The putative *Martinezella* strains and the *Rhizobium* species identified in this study are highlighted in bold. The genome of *Bradyrhizobium cajani 1010* was used as the outgroup to root the tree. A pairwise percentage identity matrix of *nodC* sequences was generated using Clustal Omega and is displayed alongside the phylogenetic tree as a heatmap. Red indicates pairwise sequence identities ≥95%, whereas lower identity values are represented by white color. Tree visualization and annotation, including the heatmap, were performed using iTOL v6.

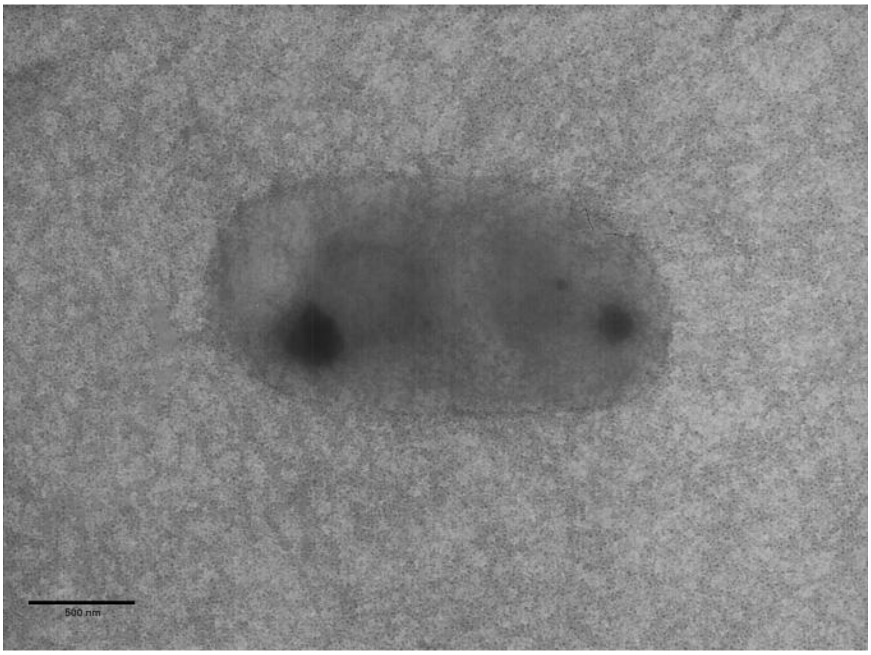


Figure S4. Transmission electron microscopy (TEM) image of *Martinezella* sp. LEGMi-135b at ×8,000.

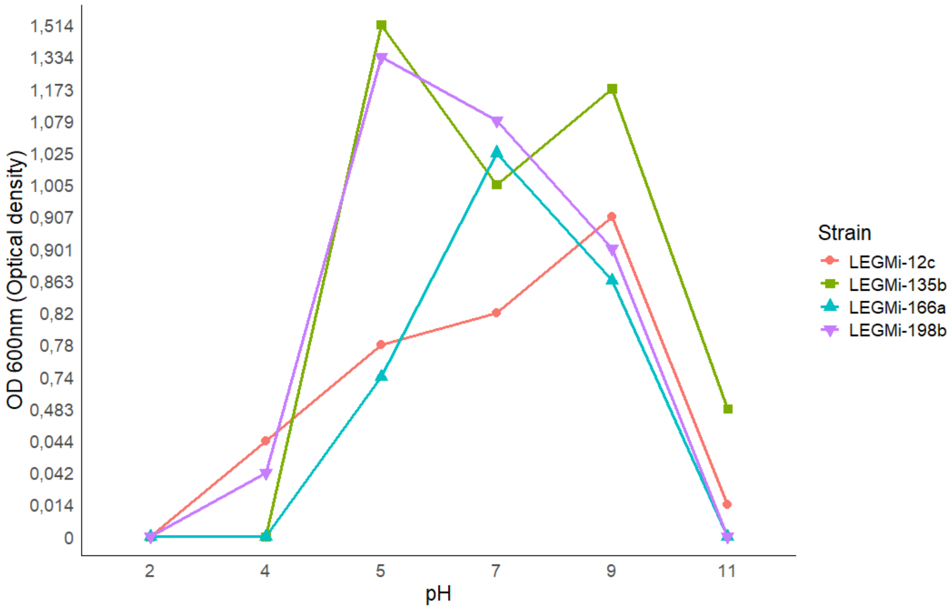


Fig. S5. Growth sensitivity of the isolated strains to different pH levels at 24h, determined via absorbance (OD 600nm).
